# Chewing-related changes in ankle muscle activity and spectral power are modulated by oral anesthesia during single-leg stance

**DOI:** 10.64898/2026.11.23.768621

**Authors:** Sebastián Jiménez-Zúñiga, Sebastian Espinoza, Wael El-Deredy

## Abstract

Mastication can facilitate lower-limb motor output, yet whether its influence on postural muscle control depends on oral sensory feedback is unknown. In a randomized, placebo-controlled, parallel-group study, 27 healthy adults performed sub-maximal single-leg stance trials at baseline, after unilateral buccal lidocaine (*n* = 12) or placebo (*n* = 15) spray, and while chewing gum. Center-of-pressure sway and surface electromyography of the tibialis anterior (TA) and medial gastrocnemius (GM) were analyzed during the first 3 s of unipedal support. Sway area showed no significant Group, Block, or interaction effects. Group differences emerged only in the change from spray alone to chewing: GM amplitude increased overall, whereas TA amplitude increased only with placebo and TA–GM coactivation only under anesthesia (Group-by-Block interactions). An exploratory cluster-based analysis, at the threshold of significance, showed greater chewing-related increases in TA spectral power (20–200 Hz) under anesthesia, early after stance onset. Participants with larger chewing-related TA power increases, most robustly at 40–80 Hz, showed smaller proportional changes in sway area, and greater trial-level TA power accompanied smaller mediolateral sway. These findings suggest that oral anesthesia alters how mastication engages early ankle neuromuscular control, revealing subtle adjustments not evident in sway area.

## Introduction

Postural control is a complex sensorimotor process that continuously integrates visual, vestibular, and somatosensory inputs to maintain the projection of the center of mass within the base of support^1^. Equilibrium emerges from the interplay between brainstem–spinal loops that regulate postural tone and cortical–subcortical–cerebellar networks that generate predictive (feedforward) and corrective (feedback) adjustments according to task demands^1–3^. Although force-platform posturography is widely used to quantify standing balance through center-of-pressure (CoP) trajectories^4,5^, CoP metrics represent only the net mechanical outcome of an abundant motor system^6,7^. Consequently, when sensory feedback is altered or balance demands increase, the central nervous system can keep postural excursion within stable bounds through adjustments in lower-limb muscle recruitment and co-contraction that are not necessarily reflected in global CoP measures^6^.

The stomatognathic system provides a dense source of proprioceptive and mechanoreceptive feedback^8,9^. Periodontal, jaw muscle spindle, temporomandibular joint, and oral mucosal afferents encode mandibular dynamics and occlusal loads through the trigeminal nerve, and trigeminal sensory and mesencephalic nuclei maintain reciprocal projections with the vestibular nuclei, reticular formation, and cerebellum, providing pathways capable of modulating antigravity muscle tone and whole-body posture^8^. Acute manipulations of trigeminal afference modify postural regulation: unilateral trigeminal anesthesia alters static balance and induces context-dependent postural deviations^10^, whereas intraoral mandibular repositioning reduces postural sway in patients with cervical sensorimotor dysfunction but not in healthy controls^11^. Moreover, unilateral chewing and masticatory muscle activity covary with trigeminal somatosensory acuity and plantar pressure distribution^9,12^, stomatognathic inputs may exert a greater influence on muscle tone and postural stability under unstable support conditions^13,14^, and occlusal status has been associated with postural stability in both bipedal and single-leg stance^15^.

Beyond passive sensory signaling, rhythmic mastication is a sensorimotor behavior that modulates widespread cortical, cerebellar, and brainstem networks^8,16^. Functional near-infrared spectroscopy has shown that periodontal tactile input during biting activates prefrontal networks, an effect abolished by periodontal ligament anesthesia but not by surface anesthesia alone^17^. Chewing has also been linked to increased locus coeruleus and reticular arousal activity, as inferred from task-related pupil dilation^8,18^, and has been proposed to act as an endogenous pacemaker that entrains frontocentral oscillatory dynamics, even when unilateral oral mucosal sensation is attenuated by topical anesthesia^16^. Posturally, chewing has been suggested to promote strategies characterized by greater compliance and lower energetic expenditure during unipedal and bipedal balance tasks^19^, whereas impaired masticatory function—clinically expressed as oral frailty—nearly doubles the risk of falls in adulthood^20^.

Evaluating the interaction between oral sensory reduction and mastication requires a postural task demanding enough to challenge distal neuromuscular coordination, such as the Single Leg Stand Test (SLST)^15,21^. When performed as repeated, sub-maximal trials that do not progress to task failure, the SLST allows examination of the early stabilization phase after the transition to unipedal support^22^, in which balance is maintained within a narrow stability margin and adjustments are expected to be subtle and primarily neuromuscular rather than evident as large changes in global sway. These demands are compensated mainly by the ankle musculature: the tibialis anterior (TA) and medial gastrocnemius (GM) act as key stabilizers, TA activity alone explains about one-third of CoP displacement variance, and forefoot–rearfoot frontal-plane instability selectively recruits the TA^23–25^. The central nervous system regulates ankle stability through a balance between reciprocal inhibition and agonist– antagonist coactivation^6,24^. TA–GM coactivation, quantified by indices such as the Rudolph co-contraction index^26^ or the Falconer–Winter ratio^27^, increases apparent ankle impedance as task difficulty or sensory uncertainty rises^6,23^, but carries a higher neurometabolic cost, does not necessarily improve mechanical stability, and, at least in older adults, covaries with greater instantaneous postural sway^6,28^.

Stabilization during challenging single-leg tasks involves increased activity of sensorimotor cortical areas contralateral to the stance leg^21^ and selective presetting of corticospinal excitability targeting the TA rather than the plantarflexors^29^. Whereas surface electromyography (sEMG) amplitude envelopes reflect global muscle excitation and are sensitive to processing choices^30,31^, time–frequency decomposition of nonrectified sEMG can capture spectral features associated with descending neural drive^32,33^. TA motor unit synchronization in the Piper band (30–60 Hz) is coherent with contralateral motor cortex activity^34^, cortico-muscular coupling emerges during reactive stepping^35^, and Piper-band cortico-synergy coherence during perturbed unipedal stance decreases as balance improves with practice^36^. Furthermore, intramuscular couplings between gamma-band (60–80 Hz) amplitudes and lower-frequency bands, including the Piper band, carry task-relevant information associated with lower balance error on an unstable board^37^.

Previous studies have examined trigeminal sensory perturbation^10^ or mastication^19^ in isolation, or have relied on kinematic and posturographic outcomes that can mask underlying muscular reorganization^7,15^. Consequently, it remains unknown how rhythmic mastication interacts with an acute reduction in oral mucosal afference to modulate ankle muscle excitation, agonist–antagonist coactivation, and TA time–frequency dynamics during the early stabilization phase of unipedal stance. To address this gap, we combined force-platform posturography with lower-limb sEMG amplitude, TA–GM coactivation, and instruction-locked time–frequency analyses during the SLST in healthy adults randomized to unilateral buccal local anesthesia or placebo across baseline, no-chewing (spray only), and chewing (spray plus gum) blocks. Specifically, we asked (i) whether oral sensory reduction and mastication would alter CoP sway during the early stabilization phase of the SLST; (ii) whether chewing-related changes in ankle muscle amplitude and TA–GM coactivation would depend on oral sensory input; and (iii), in exploratory analyses, whether TA time–frequency power showed group-dependent changes and whether these neuromuscular changes covaried with postural sway. We hypothesized that oral sensory reduction would modify the ankle muscle response to mastication.

## Results

### Participants and included observations

Thirty participants were recruited. Three allocated to anesthesia were excluded because of sEMG signal loss, leaving 27 participants for analysis (anesthesia, *n* = 12; placebo, *n* = 15; Supplementary Table S1). After trial selection, 1,209 observations across the three blocks were analyzed, including 807 no-chewing and chewing trials in the baseline-referenced models. Valid TA time–frequency (TF) estimates were available for 1,171 trials, including 781 no-chewing and chewing trials matched to posturography.

### No significant group or block effects were detected in baseline-referenced posturography

For the primary outcome, 95% CoP sway ellipse area, the trial-level LMM showed no significant effect of Group (*F*_1,25.02_ = 0.566, *P* = 0.459), Block (*F*_1,778.11_ = 0.018, *P* = 0.894), or their interaction (*F*_1,778.11_ = 0.352, *P* = 0.553). None of the ten secondary outcomes showed a significant Group, Block, or interaction effect after false-discovery-rate correction, and no simple contrast survived Holm correction (Supplementary Tables S2 and S3).

### GM RMS increased overall with chewing, with group-dependent TA changes

GM RMS amplitude increased overall from the no-chewing to the chewing block (*F*_1,803_ = 12.53, *P <* 0.001 for Block), without a significant Group-by-Block interaction (*F*_1,803_ = 0.180, *P* = 0.672), whereas TA RMS showed a Group-by-Block interaction (*F*_1,803_ = 13.02, *P <* 0.001). In placebo, chewing-related increases were 10.0% for TA (95% CI, 3.3 to 17.2%; *P*_Holm_ = 0.013) and 8.1% for GM (95% CI, 2.7 to 13.8%; *P*_Holm_ = 0.012; Fig. 1a,b). In anesthesia, neither change survived Holm correction (TA, *−*7.6%, *P*_Holm_ = 0.085; GM, +6.3%, *P*_Holm_ = 0.110).

**Figure 1.**
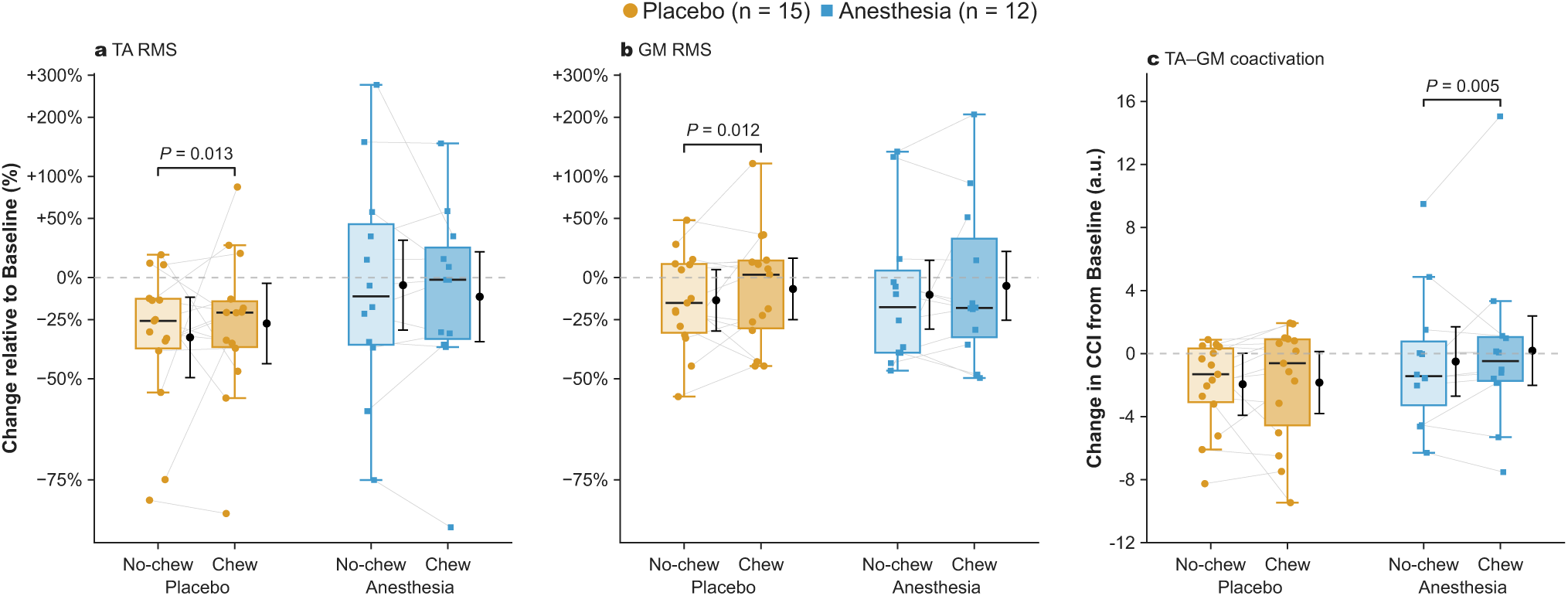
RMS amplitude increases in placebo and TA–GM coactivation increases in anesthesia from No-chew to Chew. **a,b**, Tibialis anterior (TA) and medial gastrocnemius (GM) RMS amplitude relative to each participant’s mean Baseline value, plotted on a log-ratio scale with ticks labelled as percentage change. **c**, Change in the mean Rudolph co-contraction index (CCI) from each participant’s mean Baseline value (arbitrary units). Boxes summarize participant-by-block means of the available trials in the corresponding analysis scale (placebo, *n* = 15; anesthesia, *n* = 12). Center lines indicate medians, boxes the interquartile range (IQR), and whiskers the most extreme observations within 1.5 *×* IQR below the lower or above the upper quartile. All participant values, including those beyond the whiskers, are shown as circles (placebo) or squares (anesthesia); grey lines connect No-chew and Chew values from the same participant. Black markers and vertical bars indicate trial-level linear mixed-effects model (LMM) estimates of group-by-block means and their nominal 95% confidence intervals. Brackets indicate significant within-group Chew minus No-chew contrasts; displayed *P* values are Holm-adjusted across four simple contrasts per muscle–outcome analysis. The horizontal dashed line indicates no change from Baseline. Baseline denotes the block without oral intervention; No-chew and Chew denote the assigned oral intervention without and with rhythmic mastication, respectively. RMS reflects the overall excitation of each muscle separately, whereas the CCI reflects how similar TA and GM activation levels are at each instant, weighted by their combined activity. Thus, in placebo both muscles increased in parallel without changing their balance, whereas under anesthesia TA and GM activation became more similar in magnitude, increasing coactivation without a net TA increase. RMS, root mean square.

Mean-peak amplitude showed the same pattern, with a Group-by-Block interaction for TA (*F*_1,803_ = 12.28, *P <* 0.001), an overall Block effect for GM (*F*_1,803_ = 12.44, *P <* 0.001; interaction *P* = 0.661), and chewing-related increases in placebo of 10.0% for TA and 8.2% for GM (both *P*_Holm_ *≤* 0.015). TA peak amplitude also showed a Group-by-Block interaction (*F*_1,803_ = 11.77, *P <* 0.001) and decreased with chewing in anesthesia (*−*9.8%; 95% CI, *−*16.8 to *−*2.3%; *P*_Holm_ = 0.045), whereas GM peak amplitude showed no significant effects. No anesthesia–placebo contrast within either block was significant for any amplitude outcome.

### TA–GM coactivation increased with chewing in anesthesia

Mean Rudolph CCI showed a Block effect (*F*_1,803_ = 7.69, *P* = 0.006) and a Group-by-Block interaction (*F*_1,803_ = 4.14, *P* = 0.042), increasing with chewing in anesthesia (0.687 arbitrary units; 95% CI, 0.270 to 1.105; *P*_Holm_ = 0.005) but not in placebo (0.106 units; 95% CI, *−*0.269 to 0.481; *P*_Holm_ = 0.678; Fig. 1c). No anesthesia–placebo contrast within either block was significant. The Falconer–Winter index showed the same pattern (Group-by-Block *F*_1,803_ = 16.52, *P <* 0.001; chewing-related increase in anesthesia, 4.67 percentage points; 95% CI, 2.91 to 6.42; *P*_Holm_ *<* 0.001). Simultaneous TA–GM activation increased overall with chewing (*F*_1,803_ = 8.51, *P* = 0.004 for Block) without a Group-by-Block interaction (*P* = 0.894) or significant simple contrasts, and peak Rudolph CCI and its latency showed no significant effects.

### Chewing-related TA power changes differed between groups in a permutation-defined cluster

The direct anesthesia-minus-placebo comparison of chewing-related TF changes (chewing minus no-chewing maps) identified one positive TA cluster (*P*_FWER_ = 0.0498, 5,000 permutations; Fig. 2a), indicating a greater chewing-related power change in anesthesia than placebo rather than an increase shared by all participants. This was the only cluster meeting the bilateral FWER threshold among the six muscle-by-contrast tests; neither within-group TA test nor any GM test reached significance. The cluster-level result was close to the conventional 0.05 significance threshold.

**Figure 2.**
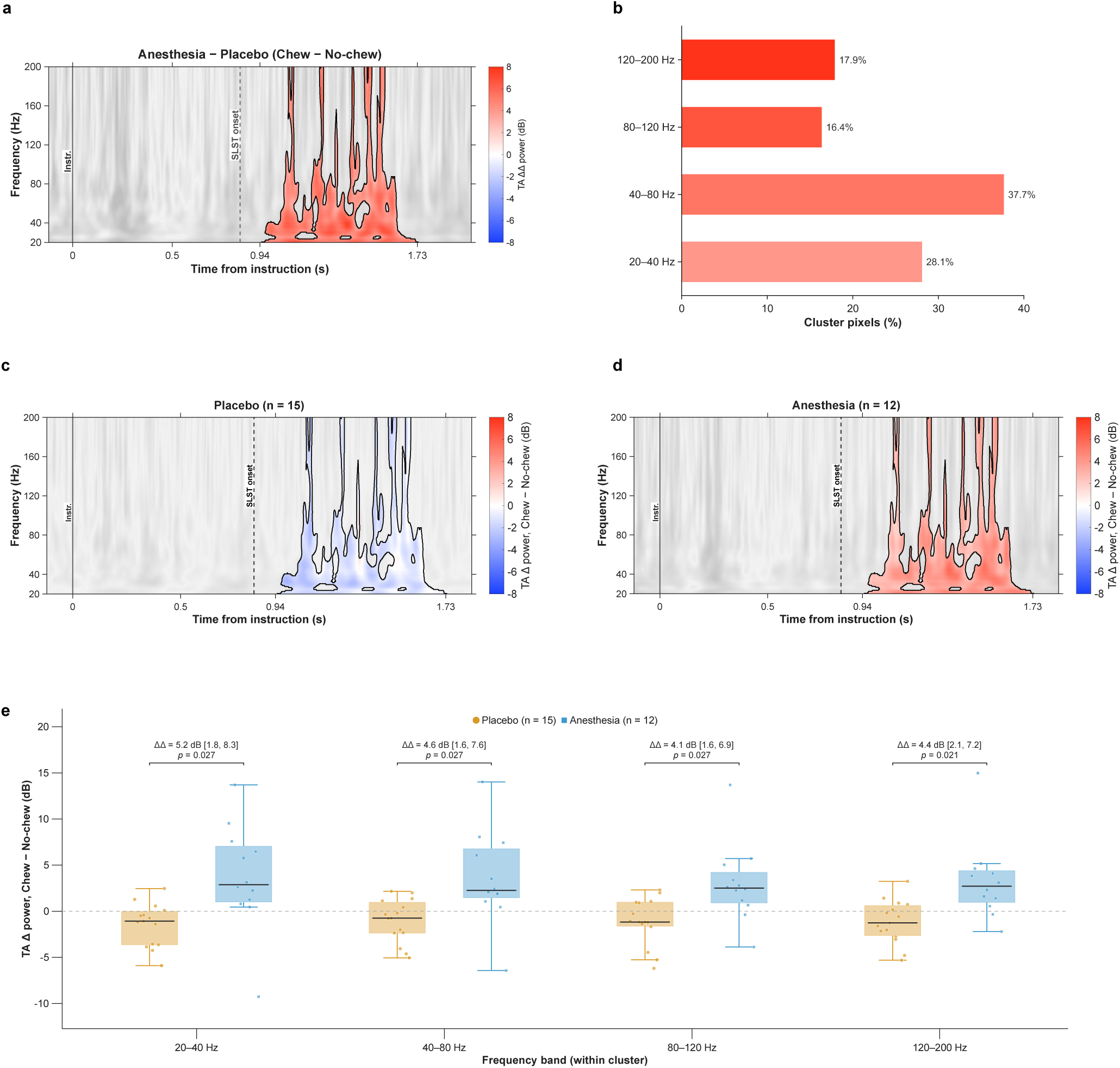
Chewing-related TA power changes are greater in anesthesia than placebo within a permutation-defined time–frequency cluster. **a**, Between-group difference in tibialis anterior (TA) chewing-related power changes (Chew minus No-chew), calculated as anesthesia minus placebo (ΔΔ). The black contour delineates the positive cluster identified by the two-sided cluster-based permutation test (5,000 permutations; *P*_FWER_ = 0.0498). **b**, Percentage of cluster pixels within each frequency band; these proportions describe cluster geometry, not power contribution. **c,d**, Mean participant-level chewing-related power changes for placebo (*n* = 15) and anesthesia (*n* = 12), respectively, shown on a common color scale (*−*8 to +8 dB); **a** uses its own symmetric scale for the between-group difference. All maps are instruction-locked, with color restricted to the exact between-group cluster mask and grayscale outside it. The identical contours in **c,d** denote this shared mask, not independently significant within-group clusters. The solid vertical line marks the instruction (Instr.); the dashed line marks median SLST onset (0.840 s). A post hoc analysis realigned to each trial’s SLST onset is shown in Supplementary Fig. S1. **e**, Participant-level mean chewing-related power changes within each frequency band’s intersection with the cluster. Circles represent placebo and squares anesthesia. Center lines indicate medians, boxes the interquartile range (IQR), and whiskers the most extreme observations within 1.5 *×* IQR of the corresponding quartile. All participant values are shown. Annotations report anesthesia-minus-placebo mean differences (ΔΔ, dB), nominal 95% percentile bootstrap confidence intervals from 5,000 within-group participant resamples, and two-sided Welch *P* values Holm-adjusted across four bands. The horizontal dashed line denotes zero chewing-related change. Band-wise comparisons characterize the selected ROI in the same sample and do not independently confirm the cluster-level result. In practical terms, when chewing was added, TA power within this early time–frequency region tended to decrease in placebo and to increase under anesthesia. No-chew and Chew denote the assigned oral intervention without and with rhythmic mastication, respectively. SLST, Single Leg Stand Test; ROI, region of interest.

The nonrectangular cluster contained 45,873 time–frequency pixels and spanned 0.948–1.734 s after the instruction and 20–200 Hz, with a peak statistic of *t* = 3.75 at 78 Hz and 1.502 s. Median SLST onset occurred at 0.840 s. This exact mask defined the ROI for all subsequent analyses. The largest share of cluster pixels fell within 40–80 Hz (37.7%), followed by 20–40 Hz (28.1%), 120–200 Hz (17.9%), and 80–120 Hz (16.4%; Fig. 2b). Within the shared mask, chewing-related changes were predominantly negative in placebo and positive in anesthesia (Fig. 2c,d).

In a post hoc analysis realigned to each trial’s SLST onset, the positive between-group cluster re-emerged in the same time–frequency region (0.38–0.72 s after onset, corresponding to the time window of the original cluster relative to median onset; 20–200 Hz), with the same direction of group change, although it did not reach significance (*P*_FWER_ = 0.087; Supplementary Fig. S1). A second, late cluster (2.30–3.00 s; *P*_FWER_ = 0.069) reached the end of the analyzed window.

### Band-wise power changes within the TA cluster were greater in anesthesia than placebo

For 20–40, 40–80, 80–120, and 120–200 Hz, respectively, mean participant-level chewing-related changes within the cluster were *−*1.47, *−*1.03, *−*1.11, and *−*1.15 dB in placebo, compared with +3.69, +3.54, +2.96, and +3.26 dB in anesthesia (Fig. 2e). Within this selected ROI, anesthesia-minus-placebo differences were 5.2 dB for 20–40 Hz (95% CI, 1.8 to 8.3; *P*_Holm_ = 0.027), 4.6 dB for 40–80 Hz (95% CI, 1.6 to 7.6; *P*_Holm_ = 0.027), 4.1 dB for 80–120 Hz (95% CI, 1.6 to 6.9; *P*_Holm_ = 0.027), and 4.4 dB for 120–200 Hz (95% CI, 2.1 to 7.2; *P*_Holm_ = 0.021). Studentized permutation tests gave similar results.

### Greater ankle-muscle activity and co-contraction were associated with greater within-block sway area

Within blocks, a one-SD higher trial-level predictor value was associated with greater sway area for TA RMS (18.7%; 95% CI, 14.4 to 23.0%), GM RMS (15.4%; 95% CI, 11.2 to 19.7%), and mean Rudolph CCI (15.0%; 95% CI, 10.8 to 19.3%; all *q <* 0.001; Fig. 3a–c). Models using absolute sway-area differences from baseline also showed positive associations. In placebo, the CCI–sway association strengthened with chewing (Δ*β* = 0.127 log-area units per SD; 95% CI, 0.034 to 0.221; *P*_Holm_ = 0.045; Fig. 3c), whereas no TA or GM slope change survived Holm correction, and no between-group difference in slope change survived FDR correction (all *q ≥* 0.129).

**Figure 3.**
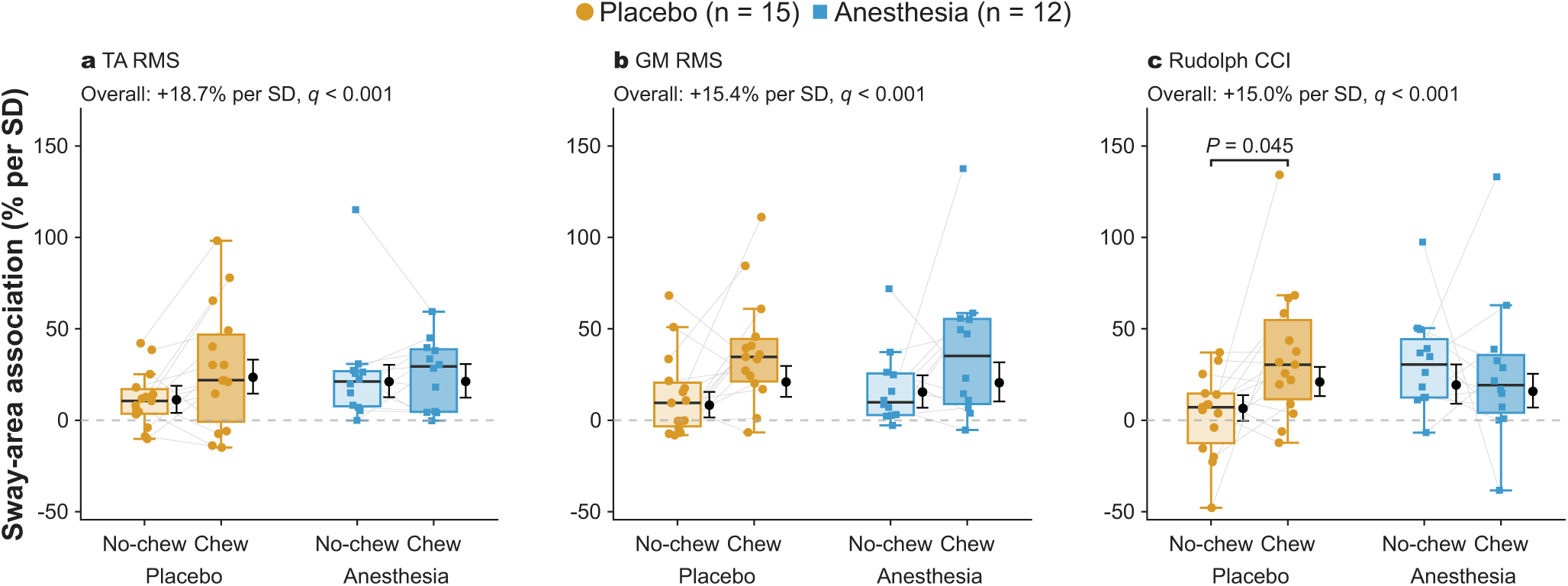
Greater ankle-muscle activity and co-contraction are associated with greater within-block sway area. Within-block associations between 95% center-of-pressure (CoP) sway ellipse area and **a**, tibialis anterior (TA) RMS; **b**, medial gastrocnemius (GM) RMS; and **c**, mean Rudolph co-contraction index (CCI). Sway area was analyzed as the natural-log ratio to each participant’s arithmetic mean Baseline value. Each point represents a participant-specific ordinary least-squares slope fitted to that participant’s trials in one block, expressed as the estimated percentage difference in sway area per one-SD higher predictor value. Predictors were centered within participant and block and scaled by a single pooled within-participant-block SD per predictor. Circles indicate placebo (*n* = 15) and squares anesthesia (*n* = 12); grey lines connect the same participant across No-chew and Chew. Boxes show medians and interquartile ranges, with whiskers extending to 1.5*×* the interquartile range; all participant values are shown. Black markers and vertical bars indicate group-by-block slopes and nominal 95% confidence intervals from trial-level linear mixed-effects models (LMMs) containing the predictor-by-Group-by-Block interaction and a participant random intercept. Slopes and confidence limits were transformed as 100[exp(*β*) *−* 1]. The “Overall” annotations report a single common slope per predictor from separate LMMs including Group, Block, and their interaction as fixed effects and a participant random intercept, fitted to all 807 No-chew and Chew trials from 27 participants. The corresponding *q* values were adjusted by Benjamini–Hochberg FDR across the three predictors; they are not tests of individual groups or blocks. The bracket in **c** identifies the chewing-related strengthening of the CCI–sway association within placebo (*P*_Holm_ = 0.045), adjusted across six slope-change contrasts (three predictors by two groups). The dashed line denotes zero association, not no change from Baseline. In practical terms, trials with higher muscle activity or coactivation showed larger sway, which is compatible with greater corrective effort rather than steadier stance. No-chew and Chew denote the oral intervention without and with rhythmic mastication; RMS, root mean square; SD, standard deviation.

### Trial-level TA TF power was associated with mediolateral sway but not sway area

Across 781 matched no-chewing and chewing trials, whole-ROI TA power was not associated with sway area (0.70% per SD; 95% CI, *−*3.23 to 4.79%; *P* = 0.731), nor was band-specific power (all *q ≥* 0.621). Greater whole-ROI power was associated with lower mediolateral RMS (*−*5.1% per SD; 95% CI, *−*7.5 to *−*2.7%; *q <* 0.001) and mediolateral range (*−*4.0%; 95% CI, *−*6.6 to *−*1.5%; *q* = 0.011), and no other secondary outcome survived correction. These mediolateral associations were negative in all four bands (RMS, *−*5.4 to *−*3.5%; range, *−*4.3 to *−*2.6% per SD; all *q <* 0.05).

### Greater chewing-related increases in TA power were associated with smaller proportional sway-area changes

At the participant level (*n* = 27), a one-SD greater chewing-related increase in whole-ROI TA power (2.83 dB) was associated with an 11.4% lower chewing/no-chewing geometric-mean sway-area ratio (95% CI, 1.7–20.1% lower; *P* = 0.024; Fig. 4a,b). Among the four frequency bands, only 40–80 Hz survived BH–FDR correction, with a 12.3% lower area ratio per SD of power increase (2.91 dB; 95% CI, 3.4–20.5% lower; *P* = 0.010, *q* = 0.040; Fig. 4b,d). Estimates for the other bands were also negative (*−*10.6 to *−*8.4% per SD) but did not survive correction (*q* = 0.070–0.116; Fig. 4b,c,e,f). Robust-regression sensitivity analyses gave similar estimates. The 40–80-Hz power change was not significantly associated with the change in mean Rudolph CCI (0.946 CCI units per SD; 95% CI, *−*0.320 to 2.212; *P* = 0.136).

**Figure 4.**
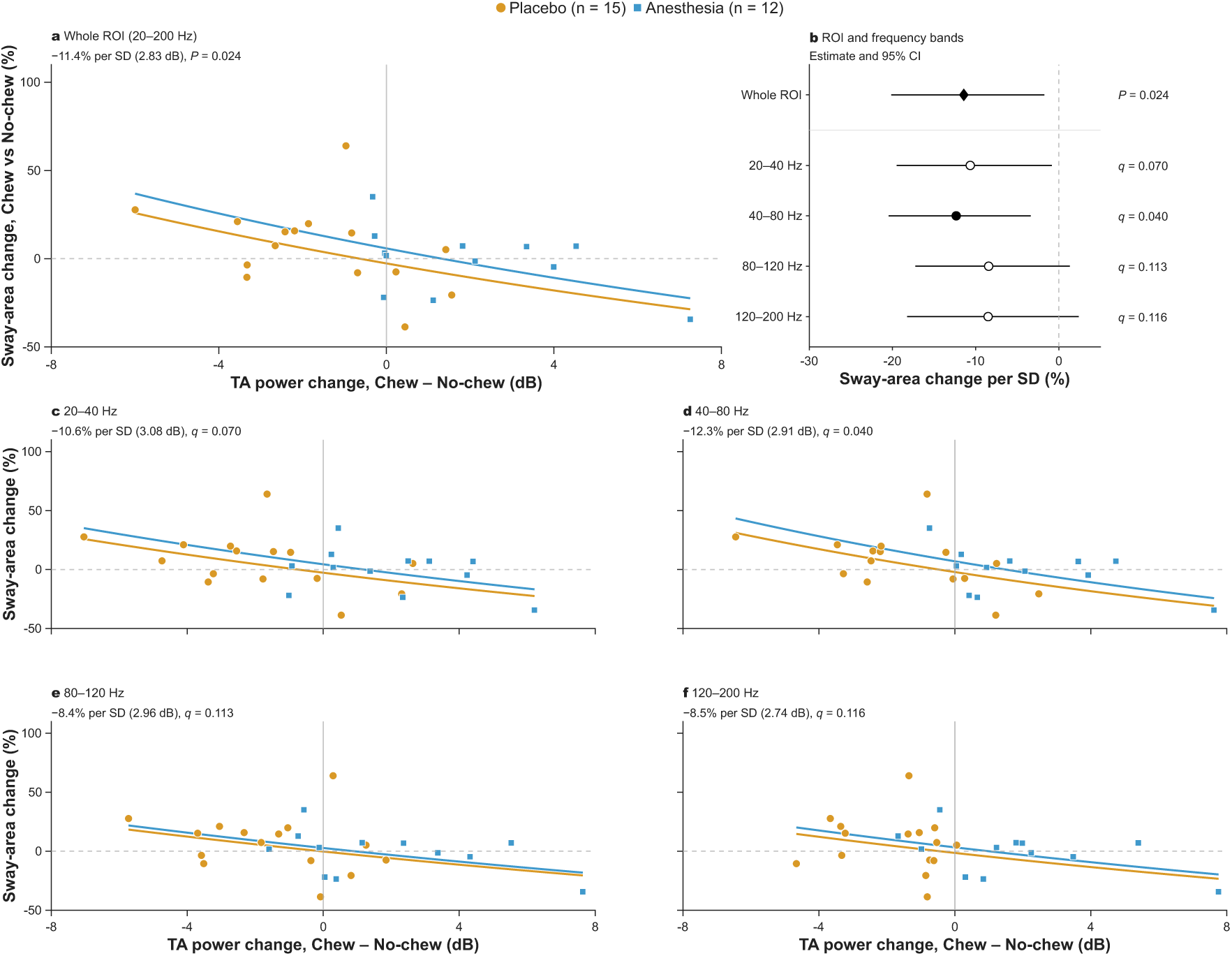
Greater chewing-related increases in TA power covary with smaller proportional sway-area changes. **a**, Association within the complete permutation-defined tibialis anterior (TA) time–frequency region of interest (ROI; 20–200 Hz). **c–f**, Corresponding associations within the 20–40, 40–80, 80–120, and 120–200 Hz portions of the same mask. Each point represents one participant (placebo, *n* = 15, circles; anesthesia, *n* = 12, squares). Within each participant and block, trial-level log sway area and TF power in decibels were averaged over the same valid trials before calculating Chew minus No-chew differences (7<u>81 m</u>atched trials overall); Baseline was not used as a reference for these changes. Displayed sway-area changes are 100[exp(Δln *A*) *−* 1], representing percentage changes in geometric-mean 95% center-of-pressure sway ellipse area. Colored curves are back-transformed predictions from group-adjusted ordinary least-squares models, with a common power-change slope on the log scale and group-specific intercepts. **b**, Forest plot of model coefficients and nominal 95% confidence intervals, transformed as 100[exp(*β*) *−* 1]. Estimates express the percentage difference in the Chew/No-chew geometric-mean area ratio per one-SD higher power change; each region uses its own between-participant SD. The diamond denotes the complete ROI and circles denote frequency bands. Filled symbols indicate nominal *P <* 0.05 for the complete ROI or BH–FDR-adjusted *q <* 0.05 for a band; open symbols indicate bands not meeting that threshold. FDR correction covered the four band tests, not the complete-ROI test. Only 40–80 Hz survived this correction. Horizontal dashed lines in the scatter panels denote zero sway-area change; vertical reference lines denote zero power change in the scatter panels and zero association in the forest plot. Band-wise findings characterize the ROI selected in the same participants and are not independent confirmation of its selection. In practical terms, participants whose TA power increased more when chewing was added showed smaller increases, or larger decreases, in sway area during Chew than during No-chew. This describes differences between participants, not a group-level reduction in sway or a causal effect of mastication. No-chew and Chew denote the oral intervention without and with rhythmic mastication; SD, standard deviation.

### Within-block TF–sway slopes did not change significantly with chewing

The within-block TF–area association did not change significantly with chewing (Δ*β* = 0.026 log-area units per SD; 95% CI, *−*0.057 to 0.109; *P* = 0.536), and neither within-group slope change nor their between-group difference survived Holm correction (all adjusted *P ≥* 0.134).

## Discussion

Baseline-referenced posturography showed no significant Group, Block, or interaction effects, yet the transition to mastication was accompanied by reorganizations in ankle-muscle sEMG amplitude, TA–GM coactivation, and TA time-frequency (TF) power, with Group-by-Block interactions for specific neuromuscular outcomes. Muscle–sway associations depended on the outcome domain and analytical level: conventional sEMG amplitude and coactivation covaried positively with sway area within blocks, TA TF power covaried negatively with mediolateral sway at the trial level, and participant-level TF power increases— most notably within the 40–80 Hz band—covaried with smaller proportional chewing-related sway-area changes. Together, these findings suggest that mastication acted as a modulator through which oral sensory input shaped ankle neuromuscular control, expressed as subtle modulations within the early stabilization phase of a sub-maximal SLST rather than as large changes in global sway.

The posturographic findings should not be read as an absence of postural regulation changes, because the models neither detected differences nor established formal equivalence with baseline. Earlier investigations reported reduced CoP excursion during gum chewing in quiet bipedal standing^38^ and decreased center-of-gravity velocity during jaw clenching on a foam surface^39^. Those bipedal paradigms differ from the early 3-s window examined here: transferring the center of gravity onto a unipedal base constitutes a demanding point-to-point control task governed by speed–accuracy trade-offs, in which the initial seconds are dominated by rapid dynamic braking and frontal-plane stabilization rather than stationary quiet standing^22^. Moreover, the SLST was performed as repeated, sub-maximal trials that did not progress to task failure; within this design, large changes in sway were not expected, and the relevant information lies instead in subtle neuromuscular modulations that maintain balance within a narrow stability margin. Consistent with this, recent work comparing chewing and handgrip conditioning prior to unipedal balance training observed divergent trajectories across area- and velocity-based CoP metrics rather than a uniform reduction in sway^19^, and jaw clenching or tongue pressing did not alter center-of-mass sway or its control on an oscillating platform^7^. Because CoP variables represent only the net outcome of redundant motor degrees of freedom^6,7^, the absence of significant CoP effects alongside sEMG modulations is compatible with participants buffering the demands of the oral task through distal neuromuscular adjustments.

The conventional sEMG amplitude results distinguish a shared, mastication-associated facilitation of the plantarflexors from a group-dependent modulation of the dorsiflexors. GM RMS and mean-peak amplitude increased overall with chewing without significant Group-by-Block interactions; although adjusted within-group GM contrasts reached significance only in placebo, the absence of an interaction precludes claiming a placebo-specific GM effect. This generalized facilitation aligns with evidence that voluntary teeth clenching induces a Jendrassik-like remote facilitation of ankle extensor and flexor spinal motoneurons^40^ that persists after moderate soleus fatigue^41^. Beyond general arousal mediated by reticular activating and locus coeruleus-noradrenergic pathways^8,18^, human reticulospinal drive shows a preferential bias toward lower-limb extensor (antigravity) muscles such as the GM^42^, facilitating their rapid recruitment^43^. In contrast, TA amplitudes showed significant Group-by-Block interactions: the placebo group increased TA RMS and mean-peak excitation during mastication, whereas TA peak amplitude decreased in the anesthesia group (Fig. 1a,b). Because moderate, balanced occlusal loading optimizes ankle dorsiflexion performance and left–right occlusal asymmetries produce asymmetrical remote facilitation in lower-limb muscles^44^, the divergent TA response may reflect the combination of unilateral mucosal sensory attenuation with lateralized chewing input, altering the scaling of peak dorsiflexor bursts rather than abolishing remote facilitation. Given normalization to the pre-instruction interval and the baseline block, these effects reflect relative task-evoked excitation rather than absolute muscle force^30^.

Coactivation indices provided a complementary view. The mean Rudolph index showed a significant Group-by-Block interaction and an adjusted chewing-related increase in the anesthesia group, with a concordant Falconer–Winter pattern (Fig. 1c)^26,27^; this convergence does not represent independent replication, because both metrics derive from the same sEMG envelopes. Binary temporal overlap, which quantifies suprathreshold activation duration rather than relative amplitude, showed only an overall Block effect. Together with evidence that voluntary teeth clenching attenuates reciprocal Ia inhibition from pretibial muscles onto the soleus^40^, this suggests a shared mastication-associated propensity for simultaneous ankle muscle firing, whereas the group-specific Rudolph increase reflects altered amplitude balancing between TA and GM under anesthesia. When sensory feedback is congruent and occlusal loading is moderate^44^, ankle control can be regulated through reciprocal inhibition and sequential agonist–antagonist adjustments^24^. Conversely, when postural demands interact with altered somatosensory feedback, Ia presynaptic inhibition of soleus afferents increases^45^, peripheral afferent deficits reduce spinal reflex excitability^46^, and agonist–antagonist common inputs increase at low frequencies attributed to subcortical sources^47^, a pattern that may increase apparent ankle impedance^6,23^. Higher excitation and coactivation did not, however, equate to quieter standing: greater trial-level TA RMS, GM RMS, and Rudolph CCI were associated with larger within-block sway areas (Fig. 3), mirroring observations, mainly in adolescents and older adults, that elevated ankle co-contraction, heightened Ia presynaptic inhibition, and higher sEMG amplitudes coincide with increased CoP and center-of-mass fluctuations^23,28,45^. Because coactivation in a free-origin kinematic chain does not mechanically anchor the body^6,28^, and stiffness and metabolic cost were not measured, these associations are compatible with greater corrective activity rather than superior coordination.

Beyond envelope amplitudes, the TF analysis revealed a group-dependent spectral modulation in the TA. The anesthesia- minus-placebo comparison of chewing-related TF changes identified a positive cluster at the threshold of significance (Fig. 2a), beginning shortly after the median SLST onset and spanning the early transition into unipedal support, whereas neither within-group TA tests nor any GM tests reached significance. After post hoc realignment to SLST onset, the cluster re-emerged in the same time–frequency region with the same direction, although without reaching significance (Supplementary Fig. S1); the following interpretation therefore remains tentative. Detection of this cluster in the TA but not the GM, although not tested by a direct TA–GM contrast, is consistent with evidence that the TA receives oscillatory corticospinal drive in the Piper range^34^ and anticipatory corticospinal presetting during postural tasks^29^, that single-leg stance engages sensorimotor cortex^21^, and that extensors such as the GM receive comparatively stronger reticulospinal drive^42^. Wavelet-based analyses of balance learning show that, as a dynamic balance task becomes more efficient with practice across days, ankle muscle sEMG intensity decreases and shifts toward lower frequency components^48^. Within this framework, the predominantly negative chewing-related power changes in placebo are compatible with progressive task economization across repeated blocks (although fatigue cannot be excluded), whereas the anesthesia group exhibited positive power shifts (Fig. 2c,d). Because rhythmic mastication entrains frontocentral cortical oscillations even under topical oral anesthesia^16^, combining asymmetric oral sensory reduction with the mechanical demands of unipedal stance^15,25^ may have prevented this spectral economization and altered descending drive to the TA, although the present data cannot identify the origin of this drive. Moreover, because sEMG spectral content also depends on muscle fiber conduction velocity, volume conduction, and tissue filtering^49^, and cortico-muscular coherence was not computed^33^, 40–80 Hz power should be interpreted as a high-frequency neuromuscular modulation rather than a direct readout of cortical gamma synchronization.

In contrast to amplitude, TF power was inversely associated with sway. At the trial level, greater TA TF power within the ROI covaried with smaller mediolateral CoP RMS and range, without a corresponding association with ellipse area. This directional specificity aligns with the functional biomechanics of the double-to-single-leg transition^22^: during unipedal stance and mediolateral perturbations, frontal-plane instability and subtalar pronation–supination shifts are counteracted in part by TA activation, which acts as an ankle invertor that can stabilize the medial longitudinal arch and constrain mediolateral CoP excursions^25,47^. Because directional RMS/range and two-dimensional prediction ellipses capture distinct geometric properties of the statokinesigram^4,5,50^, trial-by-trial increases in broad-spectrum TA drive may selectively constrain frontal-plane oscillations without altering overall planar dispersion.

At the participant level, larger chewing-related increases in TA TF power were associated with smaller proportional changes in sway area; the 40–80 Hz band was the only frequency subdivision whose association survived FDR correction, and robust regression yielded estimates of similar direction and magnitude (Fig. 4). These associations describe inter-individual differences rather than uniform reductions in sway and do not establish causality. The divergence between trial-level and participant-level associations reflects complementary analytical scales rather than a statistical contradiction: centering predictors within clusters separates within-cluster associations from between-cluster differences, which would otherwise be conflated^51^. The absence of a significant association between 40–80 Hz power changes and CCI changes is compatible with evidence that, during standing, agonist–antagonist common inputs increase mainly at low frequencies attributed to subcortical sources, whereas agonist–agonist common inputs increase in the corticospinal beta range^47^. Thus, high-frequency TA modulation and agonist–antagonist coactivation may represent partly distinct neuromuscular adjustments—one linked to greater coactivation and instantaneous sway, and the other associated with smaller proportional sway-area changes.

Notably, no anesthesia–placebo contrast was significant in the no-chewing block; group differences emerged as Group-by- Block interactions in the transition to chewing rather than as between-group differences within a single block. This suggests that mastication—an oral motor task that depends on continuous oral feedback—acted as a sensorimotor challenge that revealed the contribution of oral sensory input to distal postural control. The following mechanisms remain working hypotheses. Unilateral buccal anesthesia attenuates superficial trigeminal mucosal afference without blocking deep periodontal ligament mechanoreceptors, temporomandibular joint receptors, jaw-closing muscle spindles, or the brainstem central pattern generator driving mastication^8,16,17^. Primary afferents from jaw-closing muscle spindles and many periodontal mechanoreceptors have their cell bodies in the trigeminal mesencephalic nucleus (Me5), which integrates jaw proprioception with the control of rhythmic elevator-muscle loading during mastication^52,53^. When unilateral mucosal anesthesia is combined with right-sided gum chewing, the resulting discrepancy between attenuated superficial mucosal feedback and intact deep Me5 proprioceptive drive—compounded by laterally unbalanced stomatognathic input^44^—may generate an orofacial sensorimotor mismatch or prediction error. Much like peripheral afferent disruptions in the lower limb alter spinal reflex gain and demand compensatory supraspinal reorganization^46,54^, this trigeminal sensory incongruence may prompt central sensory reweighting, shift attentional resource allocation, or modify descending reticulospinal and corticospinal drive during challenging balance^2,10,55^. However, because tactile thresholds were not quantitatively verified throughout the blocks and masticatory muscle sEMG or chewing kinematics were not recorded during self-paced gum chewing^9,12,44^, we cannot exclude the possibility that subtle between-group differences in jaw motor behavior or bite force contributed to the observed lower-limb responses.

Methodological strengths of this work include synchronized multimodal acquisition, rigorous trial-level quality control, preservation of trial identity, and mixed-effects modeling with serial-correlation and residual diagnostics that leveraged repeated observations while separating within-block trial covariation from block-level means^56^. Matching valid trial sets across TF and posturographic pipelines, retaining the exact nonrectangular ROI geometry, and explicitly distinguishing common-slope from slope-change models ensured analytical transparency.

Nevertheless, important limitations temper our conclusions. First, the sample fell below the initial target, and the power calculation was based on a paired comparison rather than the mixed-effects models implemented. All three complete-recording exclusions occurred in the anesthesia group, and an age imbalance existed between groups (32.7 *±* 7.1 vs. 24.1 *±* 5.9 years); because age can influence TA muscle fiber conduction velocity dispersion^57^, intracortical inhibition^58^, and beta-gamma oscillatory drive^59^, this imbalance limits the certainty of between-group inferences. Temporomandibular disorders were not formally screened, so subclinical dysfunction could have influenced masticatory behavior despite no reported pain or difficulty. Second, the block order was fixed: the chewing block was always performed last, with no additional spray-only block for comparison. Because both groups followed the same sequence, learning or fatigue are unlikely to explain the Group-by-Block interactions unless they differed between groups (e.g., because of the age imbalance); however, within-group chewing-related changes, such as the overall GM increase or the negative TF power changes in placebo, cannot be attributed specifically to mastication rather than to temporal order, learning, or fatigue^19^. Notably, the increased high-frequency TA power in the anesthesia group runs opposite to the shift toward lower frequencies expected with prolonged lower-limb activity^60,61^, and jaw-clenching remote facilitation of ankle motoneurons persists despite moderate plantarflexor fatigue^41^. Third, the 3-s windows do not characterize sustained unipedal stance^22^. Finally, the lack of preregistration, exploration of cluster settings before the reported configuration, the marginal cluster *P* value (*P* = 0.0498), which re-emerged in the same region but was not significant after realignment to SLST onset (*P* = 0.087), and reuse of a data-derived ROI for secondary regressions mean that the spectral findings are exploratory until replicated with a prospectively defined ROI in an independent cohort; cluster inference also does not establish the significance of individual pixels or exact time–frequency limits.

Future investigations should employ counterbalanced or crossover block designs with larger, age-balanced cohorts, in- corporate quantitative trigeminal sensory testing alongside bilateral masseter/temporalis sEMG and occlusal balance moni-toring^9,12,44,53^, and integrate ankle kinematics with mobile brain–body imaging (EEG–EMG coherence)^16,21,35,55^. Within the boundaries of the present study, we conclude that the transition to rhythmic mastication during early single-leg stance was accompanied by subtle ankle neuromuscular changes that differed between oral anesthesia and placebo: TA facilitation with placebo, and increased TA–GM coactivation under anesthesia, together with a tentative early increase in TA spectral power. Sway area showed no significant Group, Block, or interaction effects, whereas participants with larger chewing-related increases in TA spectral power—most robustly in the 40–80 Hz band—showed smaller proportional changes in sway area. Given that oral frailty is associated with an increased risk of falls^20^, these findings support further investigation of how oral sensorimotor function contributes to lower-limb postural control.

## Methods

### Participants and experimental design

Healthy adults of both sexes aged 18–40 years were recruited by convenience sampling from the university community of Universidad de Valparaíso. The study was conducted between March and October 2025. Eligibility required unassisted single-leg stance and right-hand dominance. All participants also reported right-leg dominance, defined as the preferred leg for spontaneously kicking a ball. Exclusions included neurological, musculoskeletal, or vestibular disorders, uncorrected visual impairment, use of psychotropic medication or other medications affecting balance or motor control, acute pain, or recent lower-limb injury. The principal investigator randomized allocation to oral anesthesia or placebo 1:1 using Research Randomizer 4.0^62^. Similar bottles were numbered 1 or 2. Participants were not informed of their assigned spray, and the acquisition technician was blinded. A different person administered the spray and knew the assignment. Thirty participants were enrolled. Three participants allocated to anesthesia were excluded from all analyses because of surface electromyography (sEMG) electrode signal loss, including posturography despite available recordings, to retain a common cohort for unimodal and multimodal analyses. The analyzed sample comprised 27 participants (anesthesia, *n* = 12; placebo, *n* = 15). Participant characteristics and exclusions are summarized in Supplementary Table S1.

An initial G*Power 3.1^63^ calculation for a paired-samples comparison (*d_z_* = 0.5, two-sided *α* = 0.05, 80% power) indicated 34 participants, with a recruitment target of 37 allowing for attrition or unusable recordings. The analyzed sample fell below this target. This calculation was not specific to the Group-by-Block interaction or association models.

The randomized parallel-group study used three fixed-sequence blocks of the Single Leg Stand Test (SLST; Supplementary Fig. S2): a baseline block without oral intervention, a no-chewing block (spray only), and a chewing block (spray plus gum). Before the no-chewing block, two puffs of commercial Xylonor Spray (Septodont, Saint-Maur-des-Fossés, France; lidocaine 150 mg/g and cetrimonium 1.5 mg/g) or placebo were applied to the left buccal mucosa, followed by 5 min of rest. Each active puff delivered approximately 10 mg of lidocaine and 0.1 mg of cetrimonium, totaling approximately 20 mg and 0.2 mg per application, respectively, according to the manufacturer’s product information^64^. The placebo was prepared by personnel not involved in the experiment, contained no lidocaine, and had a similar bitter taste. The number of puffs and application timing were matched between groups. The same two-puff intervention and 5-min rest were repeated before the chewing block, which added self-paced rhythmic right-sided mastication of custom-prepared chewing gum (chewing rate was not quantified). Participants were instructed at the start of the experiment to chew on the right side. The anesthetic was applied contralateral to the chewing side to preserve protective feedback on the working side; because mandibular movement and trigeminal integration are bilateral, this was expected to attenuate rather than abolish oral somatosensory input during mastication. A topical spray reduced mucosal input without the nociceptive stimulus of an injection^16^. Mastication began at the start of the chewing block and continued throughout the block, including the intertrial rest periods, using the same gum unit. Participants were asked about unusual sensations, accidental biting, or other problems with the gum; none were reported. No formal sensory assessment was performed to verify the anesthetic effect during the study.

The chewing gum contained damar gum as the elastomer, white beeswax to adjust consistency, saccharin as a sweetener, and soy lecithin to promote cohesion. Damar gum and beeswax were melted together at 200 *^◦^*C, then saccharin and lecithin were incorporated. The mixture was poured into silicone molds, cooled to 25 *^◦^*C, and stored in sealed packaging.

Before the protocol, participants completed up to ten familiarization SLST repetitions. A 3-min seated resting-state recording with forward gaze preceded the experimental blocks but was not analyzed. The SLST was performed barefoot on the dominant limb, with hands on the iliac crests. Each block comprised sixteen 10-s trials separated by 5 s of bipedal rest, with up to fifteen trials retained. The first trial was omitted by default; it could replace a later unusable trial if it met signal-quality and temporal-completeness criteria. Task failures, including contralateral foot contact, removal of the hands from the iliac crests, or loss of balance, were recorded for each experimental session and the affected trials were excluded during data processing. Additional trials with unusable recordings were excluded, so the number of retained trials could differ between participants and blocks. Block contrasts were feasible because all blocks used the identical SLST protocol (sixteen 10-s trials on the same limb and posture) and differed only in the oral intervention, and because sub-maximal trials in young healthy adults avoided task failure, yielding comparable trial sets across blocks and participants.

The Accredited Ethics Committee of the Faculty of Medicine, Universidad de Valparaíso approved the protocol (No. 15/2025). All procedures were conducted in accordance with relevant guidelines and regulations, including the Declaration of Helsinki. Written informed consent was obtained from all participants. The study was not preregistered.

### Acquisition, processing, and trial identification

Ground-reaction forces and moments were recorded at 1,000 Hz with a calibrated force platform (BP5046, Bertec Corporation, Columbus, OH, USA) to derive mediolateral (ML) and anteroposterior (AP) center-of-pressure (CoP) coordinates. sEMG was recorded at 4,096 Hz with a BioSemi ActiveTwo system (BioSemi B.V., Amsterdam, Netherlands) from tibialis anterior (TA) and medial gastrocnemius (GM) on the support limb and from rectus femoris (RF) on the raised limb. CMS and DRL electrodes were placed in the scalp EEG cap; EEG data were not analyzed in this study. With participants seated, the skin was cleaned with isopropyl alcohol and active surface electrodes were placed with 20-mm inter-electrode spacing following SENIAM recommendations^65^. TA electrodes were positioned approximately one-third of the distance from the fibular head to the medial malleolus; GM electrodes were placed over the most prominent part of the medial muscle belly; RF electrodes were positioned halfway between the anterior superior iliac spine and the superior patellar border. Pairs were aligned with the muscle fibers, secured with double-sided tape and additional adhesive fixation, and converted offline to bipolar derivations. RF was an auxiliary channel and was not an outcome. Acquisition used two dedicated computers: one controlled the protocol in PsychoPy 2024.2.4^66^ and generated experimental-event markers; the second recorded BioSemi sEMG, force-platform signals, and the experimental-event markers transmitted through a wired connection. These markers provided the temporal reference for signal alignment and segmentation; transmission latency was not quantified. Instructions were presented on a 144-Hz screen positioned 60 cm from participants, with its center aligned with nose height to maintain head alignment and avoid downward neck flexion.

Processing used MATLAB R2024b^67^ with the Signal Processing Toolbox (pipeline in Supplementary Fig. S3). CoP segments were detected from the ML signal using a second-order-prototype 0.05–5-Hz Butterworth band-pass filter applied forward and backward, 100-ms smoothing, and hysteresis/moving-variability onset refinement. Automatic segments began 0.30 s after the refined detection point. Detected segments were visually reviewed and selected; additional segments could be defined manually by selecting their start on the CoP trace. Each segment retained an automatic/manual flag, and participant-specific graphical records were saved for verification. Filtering was used only for detection: outcomes used the first 3 s of the unfiltered accepted segment.

BioSemi data were imported through EEGLAB^68^, resampled to 1,024 Hz, demeaned, and band-pass filtered at 20–450 Hz using a Butterworth filter with a fourth-order prototype, applied forward and backward for zero phase. Channel quality was assessed from spectra, 50-Hz signal-to-noise ratio, and kurtosis. Welch spectra used 2,048-sample Hamming windows, 50% overlap, and a 4,096-point FFT. The 50-Hz quality measure compared integrated 20–450-Hz power with 48–52-Hz power; the latter may include physiological sEMG activity. Channels were pre-specified for line-noise removal if 48–52-Hz power reached 15% of 20–450-Hz power. No channel reached this threshold, so no notch filter was applied (Supplementary Fig. S4).

A functional sEMG onset marker (SLST onset) was detected relative to each instruction using 50-ms RMS envelopes. Muscle-specific adaptive thresholds were calculated as *µ* + 4.5*σ* from the interval *−*1.5 to *−*0.5 s before the instruction. The search interval was 0.5–3 s after the instruction. The primary criterion was a simultaneous TA–GM threshold exceedance sustained for at least 150 ms; if this was not detected, the same persistence criterion was applied to TA alone. If neither criterion was met, the detector assigned a default marker at 0.9 s after the instruction rather than a detected activation onset. RF was available for graphical inspection but was not part of the quantitative detector. Detected markers were inserted into the continuous datasets, temporally ordered, and checked for internal consistency; the automatically assigned markers were retained without manual adjustment. Continuous block segments retained the internal events, extending from 3 s before the first instruction to the final trial-end marker considered (up to sixteen trials). Subsequent normalized epochs extended for 7 s from each SLST onset marker. Amplitude was quantified over 0–3 s after SLST onset. TA–GM coactivation was quantified over an equally long 3-s interval, beginning 200 ms after SLST onset (0.2–3.2 s), to characterize coactivation following the immediate onset period. Centered 50-ms RMS envelopes were computed over 0–4 s before extracting the coactivation interval, ensuring that all analyzed envelope samples used complete RMS windows without endpoint shortening. Signals were divided by the median across trials of the RMS in the 3 s before each instruction, calculated separately for each participant, block, and muscle. This signal scaling was distinct from outcome normalization to the baseline block. Amplitude log-ratios therefore compared task activity relative to each block’s own pre-instruction interval, rather than unscaled EMG amplitudes.

A canonical inventory linked modalities by participant, block, and original experimental trial number, preserving identity despite missing observations. Each modality’s analysis window was referenced to its own onset (CoP-based onset or sEMG SLST onset). It contained 1,209 trials across the three blocks, including 807 no-chewing and chewing trials. Analyses excluded observations missing required measurements without imputation.

### Posturographic and sEMG outcomes

The primary posturographic outcome was the 95% CoP sway ellipse area (cm^2^), computed for each trial, not from pooled trajectories, as^5,69^

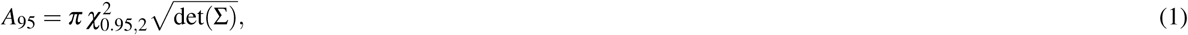

where Σ is the robust ML–AP covariance matrix estimated with MATLAB’s robustcov function (Method=’fmcd’, Fast Minimum Covariance Determinant^70^; default OutlierFraction=0.5) to limit the influence of brief outlying CoP excursions, and *χ*^2^_0.95.2_ is the 95th percentile of the chi-squared distribution with two degrees of freedom.

Ten secondary outcomes were major- and minor-axis diameters, ML and AP RMS and range, path length, net displacement, and mean and maximum radial displacement. RMS used coordinates centered on their trial mean; range was maximum minus minimum. Excursion and radial measures referenced the mean position in the first 100 ms. Path length summed successive Euclidean distances, whereas net displacement connected the initial reference to the final sample. Mean velocity, axis ratio, and eccentricity were descriptive; velocity was not tested independently of path length because all windows lasted 3 s.

For baseline-referenced analyses, positive posturographic and amplitude outcomes *X* in the no-chewing and chewing blocks were expressed as

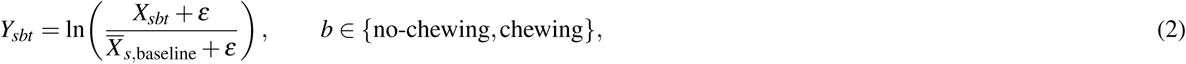

where *s*, *b*, and *t* index participant, block, and trial, *X̄_s,_*_baseline_ is the participant’s arithmetic mean baseline value, and *ε* = 10*^−^*^6^ for amplitude outcomes (*ε* = 0 for posturography). Log-scale fixed-effects estimates and confidence limits were back-transformed as 100[exp(*·*) *−* 1]. Baseline trials supplied the reference but were not response observations in these models. Descriptive absolute posturographic changes were calculated by subtracting each participant’s arithmetic mean baseline value from their arithmetic mean no-chewing or chewing value, then summarized across participants as mean *±* SD, giving each participant equal weight. These original-unit summaries and the back-transformed LMM estimates were calculated on different scales and were not direct conversions of each other.

TA and GM amplitude outcomes were RMS, peak amplitude, and mean-peak amplitude (mean of rectified samples at or above the trial’s 90th percentile). No-chewing and chewing values were expressed as in Eq. (2).

Coactivation used centered 50-ms RMS envelopes. The primary measure was the temporal mean Rudolph co-contraction index^26^:

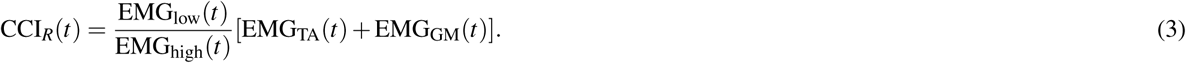

The lower/higher envelopes were identified samplewise, with zero assigned when the higher envelope was effectively zero. Peak CCI and latency were also evaluated. Simultaneous activation was the percentage of samples at which both envelopes reached at least 20% of their respective within-trial maxima. The Falconer–Winter index provided a sensitivity measure^27^. Coactivation outcomes in the no-chewing and chewing blocks were absolute differences from the mean baseline value.

### Time–frequency analysis

Time–frequency (TF) maps used filtered, nonrectified bipolar sEMG and were instruction-locked (*t* = 0). Convolution used epochs from *−*3 to 7 s to reduce edge effects. Six-cycle complex Morlet wavelets covered 20–200 Hz in 1-Hz steps at 1,024 Hz^71,72^. Each wavelet was evaluated over *−*1 to 1 s and its Fourier spectrum was normalized to unit peak magnitude. Power was the squared modulus of the convolution; maps from *−*1.5 to 4 s were retained.

Within each participant, block, and muscle, mean squared sEMG during *−*1.0 to *−*0.4 s and maximum squared sEMG during *−*1 to 4 s were standardized across trials. An absolute *z* score above 3 for either measure excluded the trial. These quality flags were intersected with the canonical inventory without recalculating thresholds, leaving 1,171 TA and 1,175 GM trials across the three blocks.

For participant-level cluster maps, linear power was averaged across valid trials (*P_sb_*) before conversion to decibels^71^

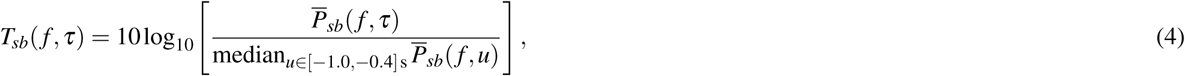

where *f* denotes frequency and *τ* denotes time relative to the instruction. Trial-level region-of-interest (ROI) extraction applied the same expression to each trial’s own power and pre-instruction reference; averaging trial-level dB values was therefore not generally equivalent to converting an averaged linear-power map. The TF reference was a pre-instruction interval within each block, not the baseline block. In the primary analysis, detected SLST onset was displayed only as a temporal reference, without realigning the maps.

### Permutation-defined ROI and frequency-band analyses

Participant maps were contrasted as *D_s_* = *T_s,_*_chewing_ *−T_s,_*_no-chewing_ (chewing-related change). Algebraically, *D_s_* = (*T_s,_*_chewing_ *− T_s,_*_baseline_) *−* (*T_s,_*_no-chewing_ *−T_s,_*_baseline_), because the same baseline map cancels. This identity does not introduce an additional baseline-block normalization. Six cluster-based permutation tests^73^ covered TA and GM within each group and between groups. Within-group tests used one-sample *t*statistics and participant sign flips; between-group tests used Welch statistics and group-label permutations preserving sample sizes. The search domain was 20–200 Hz and 0–4 s. Positive and negative clusters used *|t| >* 1.8, four-neighbor connectivity, and at least five pixels; mass summed *t* magnitudes within each same-sign cluster.

The reported analysis used 5,000 permutations, with the random-number generator seed set to 20260830 for reproducibility. Each permutation retained the maximum cluster mass across both tails, and permutation *P* values used the plus-one correction^74^. Two-sided family-wise error was controlled within each test, not jointly across the six tests. Exploratory analyses included preliminary runs with 1,000 and 10,000 permutations and alternative cluster-size settings before the reported configuration was adopted. The analysis settings were not preregistered.

After the primary analysis, a post hoc sensitivity analysis realigned each trial’s TF power to its own SLST onset (*t* = 0) before averaging within participant and block, retaining the pre-instruction reference of Eq. (4). The six cluster-based tests were repeated with the same statistics and settings over 20–200 Hz and 0–3 s after onset; the TA between-group contrast is shown in Supplementary Fig. S1. This analysis was not used to define the ROI or in any subsequent analysis.

The exact nonrectangular positive between-group TA cluster defined the ROI, rather than an arbitrarily selected window. Its enclosing limits were approximately 0.948–1.734 s and 20–200 Hz; only pixels in the mask were extracted. The original mask was fixed across groups and subsequent analyses. Cluster inference applied to the aggregate statistic, not to individual pixels or exact boundaries. ROI-based follow-up characterized this region within the same sample, conditional on its selection rather than as independent confirmation.

Mean ROI dB power was the principal trial-level TF measure, with median power as a complementary descriptor. Four nonoverlapping bands intersected the mask: 20–40, 40–80, 80–120, and 120–200 Hz, assigning shared boundaries to the higher band and including 200 Hz in the last. Cluster distribution was each band’s share of all mask pixels, summing to 100%; it described cluster geometry, not power contribution.

For band-wise group comparisons, each participant’s chewing-related dB-map difference was averaged over the band’s mask pixels. Mean anesthesia-minus-placebo changes were tested with Welch tests, Holm-adjusted across four bands. Sensitivity used 50,000 studentized Welch group-label permutations (seed 20261012), with Holm adjustment across four permutation *P* values. Figure confidence intervals used 5,000 within-group participant bootstrap resamples (seed 1), taking the 2.5th and

97.5th percentiles; Hedges’ *g* summarized standardized mean differences with small-sample correction. Confidence intervals were nominal. Holm addressed the four band tests, not prior ROI selection.

### Group and block comparisons

Trial-level linear mixed-effects models (LMMs) were fitted by restricted maximum likelihood (REML) as

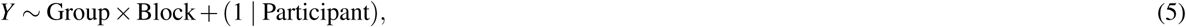

including Group, Block, their interaction, and a participant random intercept. LMMs were fitted with `fitlme` (Statistics and Machine Learning Toolbox, MATLAB R2024b); permutation and bootstrap procedures used custom code. No-chewing and chewing responses were baseline-referenced log ratios for posturography/amplitude and absolute differences for coactivation. Effects coding provided type III omnibus tests; equivalent reference coding provided simple contrasts. Posturographic inference used Satterthwaite degrees of freedom; amplitude/coactivation used residual degrees of freedom.

Four contrasts per outcome compared chewing with no-chewing within each group and anesthesia with placebo within each block, with Holm correction within each muscle–metric analysis. Amplitude/coactivation simple effects were interpreted after significant omnibus Block or interaction tests; posturographic contrasts were calculated for every metric. Area was the primary posturographic outcome, with Benjamini–Hochberg false-discovery-rate (BH–FDR) correction across the ten secondary outcomes separately for each omnibus term.

Diagnostics included Pearson-residual shape and extremes; posturographic checks also examined residual-versus-fitted and quantile–quantile plots and within-participant/block lag-one correlations. Amplitude/coactivation sensitivity fits included random Block slopes or participant-block means.

### Trial-level multimodal associations

Association LMMs used no-chewing and chewing trials and baseline-referenced log sway area as the response. Predictors were baseline-referenced log TA or GM RMS, absolute baseline-referenced mean Rudolph CCI, or trial-level mean TA ROI power in dB. Each predictor was centered within participant and block^51^ and scaled by the pooled sample standard deviation (SD) of these deviations,

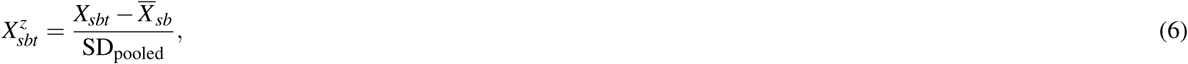

where *X̄_sb_* is the participant-by-block mean and SD_pooled_ is the single sample SD of all centered trial values (denominator *N −* 1, with *N* the number of matched trials in the model). Coefficients therefore reflect within-block, trial-to-trial associations. Separate common-slope REML models added *X^z^* to Eq. (5), with Satterthwaite inference. Each model estimated one predictor coefficient adjusted for Group, Block, their interaction, and a participant random intercept, rather than separate associations within each group or block. Coefficients were back-transformed as 100[exp(*β*) *−* 1] to express the sway-area percentage association per one pooled SD of predictor; confidence limits were transformed in the same way. These coefficients represent within-block, trial-to-trial associations, not chewing-related changes or causal effects.

The TA RMS, GM RMS, and Rudolph CCI models used 807 trials from 27 participants. Their common-slope tests formed a three-test BH–FDR family. Absolute baseline-referenced sway differences provided a sensitivity analysis. Models adding the full predictor-by-Group-by-Block interaction tested chewing-related slope changes within each group, Holm-adjusted across six contrasts (three predictors by two groups). Between-group differences in these slope changes were tested separately, with BH–FDR across the three predictors. Group-by-block LMM slopes and their nominal 95% confidence intervals were extracted from the interaction models. Participant-specific within-block OLS slopes used the same centered and scaled predictors and were displayed as descriptive boxplots and paired points; they did not supply the LMM inference.

TF and posturography were matched one-to-one by participant, block, and original trial number. Excluding 26 no-chewing and chewing observations without valid TF left 781 matched trials from all 27 participants. Whole-ROI TF association with area was evaluated at nominal *α* = 0.05, outside the ten-test BH–FDR family of secondary posturographic outcomes. Band-specific models retained the same matching and model structure, with BH–FDR across four area tests. ML RMS and ML range, selected after the whole-ROI results, formed an additional eight-test family (two outcomes by four bands). These follow-up tests were exploratory. In all trial-level TF models, sway outcomes were referenced to baseline, whereas TF power retained its pre-instruction dB reference and was centered and scaled as in Eq. (6); TF power was not normalized to the baseline block.

### Participant-level TF–sway change associations

A distinct analysis related participant-level chewing-related changes in sway and TF, using the same 781 matched trials. Trial log area was arithmetically averaged within each participant and block, and the no-chewing mean was subtracted from the chewing mean. This response was the log ratio of geometric-mean areas, not the logarithm of the ratio of arithmetic-mean areas. Trial-level ROI power in dB was averaged over the same trials before subtraction. The procedure was repeated for each of the four frequency portions of the exact ROI mask. All five analyses used the same participants and sway-area changes; each participant contributed one change pair per analysis (*n* = 27). Neither variable was normalized to baseline for these change associations.

Power changes were centered on their across-participant mean and divided by their across-participant sample SD (denominator *n−* 1), calculated separately for the complete ROI and each band. This standardization differed from the within-participant-block scaling in Eq. (6). OLS models fitted with `fitlm` regressed the log-area change on standardized power change and Group, estimating a common slope with group-specific intercepts and no power-by-Group interaction. Two-sided coefficient tests and nominal 95% confidence intervals used 24 residual degrees of freedom. The complete-ROI association used a nominal *P* value; the four band tests formed a separate BH–FDR family. No correction jointly covered the complete-ROI and band tests or the prior ROI selection.

Coefficients and their confidence limits were transformed as 100[exp(*β*) *−* 1], expressing the percentage difference in the chewing/no-chewing geometric-mean area ratio per one-SD higher power change. For scatter plots, each participant’s log-area change was transformed in the same way to display the percentage change in geometric-mean area. Colored curves were back-transformed group-specific predictions from the common-slope log-scale model, not separately fitted group slopes or estimates of arithmetic-mean area change. The forest plot showed transformed coefficients and nominal confidence intervals (Fig. 4).

The proportional models were initially implemented as sensitivity analyses and subsequently used for the participant-level presentation; they were not prospectively specified primary models. Sensitivity fits used robust regression with the same predictors and Group adjustment for the complete ROI and all four bands (`fitlm`, `RobustOpts=’on’`, bisquare weights with the default tuning constant). Their approximate *P* values were unadjusted and did not replace the OLS or BH–FDR inference. Diagnostics included residual shape, Cook’s distance, and leverage. These exploratory follow-ups were conditional on the ROI selected in the same sample.

An additional exploratory participant-level OLS model related the chewing-related change in mean Rudolph CCI to the standardized 40–80-Hz power change, adjusting for Group. Both block means used the same TF-valid trials before subtraction, without baseline normalization. The coefficient and nominal 95% confidence interval were reported in CCI units per across-participant SD of power change. This single follow-up used a nominal two-sided *P* value and was not included in the four-band TF–sway FDR family.

### Changes in within-block TF–sway slopes with chewing

A separate trial-level REML LMM added the full TF-by-Group-by-Block interaction to test chewing-related changes in the within-block association, using all 781 matched trials, baseline-referenced log sway area, the within-participant-block centered and scaled whole-ROI predictor from Eq. (6), and Satterthwaite inference. The same pooled predictor SD was retained across the no-chewing and chewing blocks. The principal contrast averaged slope changes with participant-proportion weights (15/27 placebo, 12/27 anesthesia). Three secondary contrasts—within placebo, within anesthesia, and their difference—were Holm-adjusted together. Slope changes and their confidence intervals were evaluated on the log-area-per-SD scale. Participant-specific OLS slopes were also estimated in each block using the same standardized predictor; their chewing-related differences were assessed with paired *t* and signed-rank sensitivity tests across 27 participant pairs. These tests addressed changes in within-block trial-to-trial coupling, not the association between participant-level changes described above.

All tests were two-sided at nominal *α* = 0.05. Exact *P* values, adjustment families, nominal 95% confidence intervals, and analysis-specific sample sizes were reported; there was no study-wide multiplicity correction. Group and block comparisons, within-block associations, participant-level change associations, and slope-change tests addressed distinct questions. The fixed block order meant that the chewing-related change described the transition to mastication but did not isolate mastication from time, order, or fatigue.

### Use of AI-assisted tools

Large language model tools were used to assist with code and manuscript preparation. ChatGPT (OpenAI) was used to generate and debug MATLAB code for data processing and analysis. Claude (Anthropic) was used to assist with language editing and revision of the manuscript text, generation of MATLAB code for figure preparation, and verification of reference metadata. All AI-assisted outputs were reviewed, verified, and edited by the authors. Study design, data collection, statistical analyses, and interpretation were performed and validated by the authors, who take full responsibility for the content of this manuscript.

## Code availability

The MATLAB analysis code will be made publicly available on GitHub.

## Data availability

The datasets generated and/or analysed during the current study are not publicly available because of ethics committee restrictions on participant-level data sharing, but are available from the corresponding author on reasonable request, subject to approval by the ethics committee.

## Acknowledgements

The authors thank the Faculty of Dentistry, Universidad de Valparaíso, for providing the facilities for data collection.

## Author contributions statement

S.J.-Z.: conceptualization, methodology, investigation, data curation, software, formal analysis, visualization, project adminis-tration, writing – original draft, and writing – review and editing. S.E.: conceptualization, methodology, resources (formulation and provision of the chewing gum), investigation (blinding procedure), supervision, and writing – review and editing. W.E.-D.: conceptualization, methodology, resources (equipment), supervision, and writing – review and editing. All authors interpreted the results and reviewed and approved the final manuscript.

