## Supplementary Information (Tables S1-S3, Figures S1-S4) for "Chewing-related changes in ankle muscle activity and spectral power are modulated by oral anesthesia during single-leg stance"

**Table S1.** Participant characteristics and observations included in the analyses.

| Characteristic | Anesthesia | Placebo | Overall |
|---|---|---|---|
| Participants allocated, *n* | 15 | 15 | 30 |
| Excluded because of unusable sEMG recordings, *n* | 3 | 0 | 3 |
| Participants analyzed, *n* | 12 | 15 | 27 |
| Sex, male/female, *n* | 7/5 | 9/6 | 16/11 |
| Age, years | $32.7 \pm 7.1$ | $24.1 \pm 5.9$ | $27.9 \pm 7.7$ |
| Height, cm | $172.2 \pm 8.6$ | $169.6 \pm 9.6$ | $170.7 \pm 9.1$ |
| Body mass, kg | $72.2 \pm 11.0$ | $74.1 \pm 16.6$ | $73.2 \pm 14.2$ |
| Analyzed 3-s SLST observations across the three blocks, *n* | 537 | 672 | 1,209 |
| Observations with valid TA time–frequency estimates, *n* | 516 | 655 | 1,171 |
| Individual No-chew and Chew observations included in the sEMG models, *n* | 360 | 447 | 807 |

Values are mean ± SD unless otherwise indicated. Participant characteristics describe the analyzed sample. The first trial of each block was omitted by default; it could replace a later unusable trial if it met signal-quality and temporal-completeness criteria. Additional unusable trials were excluded. Baseline, block without oral intervention; No-chew and Chew, assigned oral intervention without and with rhythmic mastication, respectively; SLST, Single Leg Stand Test; TA, tibialis anterior; sEMG, surface electromyography.

**Table S2. Posturographic changes relative to Baseline in No-chew and Chew.** Each cell combines the descriptive absolute change (mean ± SD) with the model-estimated percentage change and its nominal 95% confidence interval.

| | Placebo ($n = 15$) | | Anesthesia ($n = 12$) | |
|---|---|---|---|---|
| Outcome (absolute unit) | No-chew | Chew | No-chew | Chew |
| 95% CoP sway area[a] ($cm^2$) | $\Delta: -0.18 \pm 1.27$<br>LMM: −19.6%<br>[−28.3, −9.9]% | $\Delta: -0.06 \pm 1.01$<br>LMM: −17.3%<br>[−26.3, −7.3]% | $\Delta: -0.09 \pm 0.66$<br>LMM: −13.0%<br>[−23.4, −1.1]% | $\Delta: -0.16 \pm 1.19$<br>LMM: −14.5%<br>[−24.8, −2.9]% |
| Major-axis diameter (cm) | $\Delta: -0.17 \pm 0.50$<br>LMM: −10.1%<br>[−16.3, −3.5]% | $\Delta: -0.11 \pm 0.46$<br>LMM: −8.4%<br>[−14.7, −1.7]% | $\Delta: -0.17 \pm 0.51$<br>LMM: −9.3%<br>[−16.2, −1.8]% | $\Delta: -0.17 \pm 0.53$<br>LMM: −9.0%<br>[−15.9, −1.5]% |
| Minor-axis diameter (cm) | $\Delta: -0.03 \pm 0.28$<br>LMM: −8.8%<br>[−16.1, −0.8]% | $\Delta: -0.02 \pm 0.23$<br>LMM: −7.9%<br>[−15.3, 0.1]% | $\Delta: 0.03 \pm 0.14$<br>LMM: −2.3%<br>[−11.1, 7.3]% | $\Delta: 0.01 \pm 0.27$<br>LMM: −4.4%<br>[−12.9, 5.0]% |
| Mediolateral CoP RMS (cm) | $\Delta: 0.017 \pm 0.155$<br>LMM: −6.7%<br>[−15.8, 3.4]% | $\Delta: -0.037 \pm 0.090$<br>LMM: −11.7%<br>[−20.3, −2.2]% | $\Delta: -0.015 \pm 0.180$<br>LMM: −11.5%<br>[−21.0, −0.7]% | $\Delta: -0.037 \pm 0.142$<br>LMM: −11.1%<br>[−20.7, −0.3]% |
| Anteroposterior CoP RMS (cm) | $\Delta: -0.038 \pm 0.086$<br>LMM: −11.8%<br>[−18.2, −4.9]% | $\Delta: -0.013 \pm 0.095$<br>LMM: −8.3%<br>[−15.0, −1.1]% | $\Delta: -0.005 \pm 0.084$<br>LMM: −7.4%<br>[−14.8, 0.8]% | $\Delta: -0.053 \pm 0.107$<br>LMM: −12.8%<br>[−19.8, −5.1]% |
| Mediolateral range (cm) | $\Delta: 0.01 \pm 0.68$<br>LMM: −8.5%<br>[−18.2, 2.4]% | $\Delta: -0.16 \pm 0.50$<br>LMM: −11.9%<br>[−21.2, −1.4]% | $\Delta: -0.02 \pm 0.83$<br>LMM: −11.1%<br>[−21.6, 0.7]% | $\Delta: -0.14 \pm 0.68$<br>LMM: −11.6%<br>[−22.1, 0.2]% |
| Anteroposterior range (cm) | $\Delta: -0.20 \pm 0.40$<br>LMM: −12.8%<br>[−19.7, −5.5]% | $\Delta: -0.17 \pm 0.45$<br>LMM: −11.2%<br>[−18.2, −3.7]% | $\Delta: 0.10 \pm 0.38$<br>LMM: −4.4%<br>[−12.7, 4.7]% | $\Delta: -0.15 \pm 0.42$<br>LMM: −10.3%<br>[−18.1, −1.7]% |
| Path length (cm) | $\Delta: -0.74 \pm 1.97$<br>LMM: −4.4%<br>[−8.8, 0.3]% | $\Delta: -0.85 \pm 1.92$<br>LMM: −5.1%<br>[−9.5, −0.5]% | $\Delta: 0.36 \pm 1.77$<br>LMM: 0.6%<br>[−4.6, 6.1]% | $\Delta: -0.35 \pm 1.85$<br>LMM: −2.1%<br>[−7.2, 3.2]% |
| Net displacement (cm) | $\Delta: 0.07 \pm 0.52$<br>LMM: −16.8%<br>[−28.7, −2.9]% | $\Delta: -0.08 \pm 0.45$<br>LMM: −20.1%<br>[−31.5, −6.7]% | $\Delta: -0.02 \pm 0.73$<br>LMM: −22.4%<br>[−34.7, −7.7]% | $\Delta: -0.16 \pm 0.59$<br>LMM: −24.1%<br>[−36.1, −9.7]% |
| Mean radius (cm) | $\Delta: -0.01 \pm 0.43$<br>LMM: −12.8%<br>[−23.8, −0.3]% | $\Delta: -0.13 \pm 0.38$<br>LMM: −16.2%<br>[−26.8, −4.1]% | $\Delta: 0.00 \pm 0.63$<br>LMM: −13.9%<br>[−26.0, 0.0]% | $\Delta: -0.17 \pm 0.55$<br>LMM: −19.5%<br>[−30.7, −6.4]% |
| Maximum radius (cm) | $\Delta: -0.10 \pm 0.57$<br>LMM: −12.0%<br>[−21.0, −1.9]% | $\Delta: -0.18 \pm 0.51$<br>LMM: −12.9%<br>[−21.8, −3.0]% | $\Delta: 0.06 \pm 0.81$<br>LMM: −7.7%<br>[−18.2, 4.2]% | $\Delta: -0.19 \pm 0.69$<br>LMM: −12.8%<br>[−22.7, −1.6]% |

*First line ($\Delta$):* descriptive absolute change from Baseline, in the unit shown beside each outcome. Each participant's arithmetic mean Baseline value was subtracted from their arithmetic mean No-chew or Chew value. These participant-level differences were summarized as the group mean ± SD, giving each participant equal weight; SD denotes between-participant variability, not a standard error. *Second line (LMM):* model-estimated percentage change from Baseline. *Third line:* the corresponding nominal 95% confidence interval, also in percentage units. Trial-level LMM responses were natural-log ratios to the participant's arithmetic mean Baseline value; fixed-effects cell means and their confidence limits were back-transformed as $100[\exp(\beta) - 1]$. The absolute and percentage summaries are calculated on different scales and are not direct conversions of each other. Models included 807 No-chew and Chew trials from 27 participants, with Group, Block, their interaction, and a participant random intercept, fitted by restricted maximum likelihood. Baseline supplied the reference but was not a response block. Negative values indicate reductions relative to Baseline. Percentage confidence intervals are unadjusted for multiplicity; omnibus tests are reported in Supplementary Table S3. [a]Primary outcome. Baseline, block without oral intervention; No-chew and Chew, assigned oral intervention without and with rhythmic mastication, respectively; CoP, center of pressure; LMM, linear mixed-effects model; RMS, root mean square; SD, standard deviation.

**Table S3. Group, Block, and interaction tests for Baseline-referenced posturographic outcomes.** Omnibus tests from the trial-level No-chew and Chew linear mixed-effects models used to obtain the estimates in Supplementary Table S2.

| Outcome | Effect | $df_2$ | $F$ | $P$ | $q$ |
|---|---|---|---|---|---|
| 95% CoP sway area[a] | Group | 25.02 | 0.566 | 0.459 | — |
| | Block | 778.11 | 0.018 | 0.894 | — |
| | Group × Block | 778.11 | 0.352 | 0.553 | — |
| Major-axis diameter | Group | 25.00 | 0.001 | 0.976 | 0.990 |
| | Block | 778.08 | 0.219 | 0.640 | 0.733 |
| | Group × Block | 778.08 | 0.108 | 0.742 | 0.824 |
| Minor-axis diameter | Group | 25.02 | 0.881 | 0.357 | 0.990 |
| | Block | 778.08 | 0.053 | 0.817 | 0.817 |
| | Group × Block | 778.08 | 0.380 | 0.538 | 0.788 |
| Mediolateral CoP RMS | Group | 25.01 | 0.103 | 0.751 | 0.990 |
| | Block | 778.05 | 0.983 | 0.322 | 0.706 |
| | Group × Block | 778.05 | 1.320 | 0.251 | 0.788 |
| Anteroposterior CoP RMS | Group | 25.03 | $< 0.001$ | 0.990 | 0.990 |
| | Block | 778.11 | 0.194 | 0.660 | 0.733 |
| | Group × Block | 778.11 | 4.047 | 0.045 | 0.446 |
| Mediolateral range | Group | 25.01 | 0.029 | 0.865 | 0.990 |
| | Block | 778.05 | 0.641 | 0.423 | 0.706 |
| | Group × Block | 778.05 | 0.355 | 0.551 | 0.788 |
| Anteroposterior range | Group | 25.02 | 0.876 | 0.358 | 0.990 |
| | Block | 778.09 | 0.848 | 0.357 | 0.706 |
| | Group × Block | 778.09 | 2.833 | 0.093 | 0.464 |
| Path length | Group | 25.01 | 1.502 | 0.232 | 0.990 |
| | Block | 778.04 | 3.153 | 0.076 | 0.593 |
| | Group × Block | 778.04 | 0.961 | 0.327 | 0.788 |
| Net displacement | Group | 25.02 | 0.343 | 0.564 | 0.990 |
| | Block | 778.10 | 0.391 | 0.532 | 0.733 |
| | Group × Block | 778.10 | 0.033 | 0.856 | 0.856 |
| Mean radius | Group | 25.01 | 0.080 | 0.780 | 0.990 |
| | Block | 778.05 | 2.441 | 0.119 | 0.593 |
| | Group × Block | 778.05 | 0.159 | 0.690 | 0.824 |
| Maximum radius | Group | 25.02 | 0.110 | 0.743 | 0.990 |
| | Block | 778.06 | 1.402 | 0.237 | 0.706 |
| | Group × Block | 778.06 | 0.666 | 0.415 | 0.788 |

All models included 807 No-chew and Chew trials from 27 participants (placebo, $n = 15$; anesthesia, $n = 12$), with Group, Block, their interaction, and a participant random intercept, fitted by restricted maximum likelihood. Responses were natural-log ratios to each participant's arithmetic mean Baseline value. Omnibus tests used effects coding and the Satterthwaite approximation. Numerator degrees of freedom were 1 for every test; $df_2$ denotes denominator degrees of freedom. $P$ values are unadjusted. For the ten secondary outcomes, $q$ values apply Benjamini–Hochberg false-discovery-rate correction separately for each omnibus effect. [a]Sway area was the primary outcome, evaluated at nominal $\alpha = 0.05$ and excluded from these correction families; its $q$ values are not applicable (—). No secondary omnibus effect survived correction. Four simple contrasts per outcome compared Chew with No-chew within each group and anesthesia with placebo within each block; none was significant after Holm correction within outcome (minimum adjusted $P = 0.258$). Baseline supplied the reference but was not a response block; the Block tests compare No-chew with Chew, not Baseline with either block. CoP, center of pressure; RMS, root mean square.

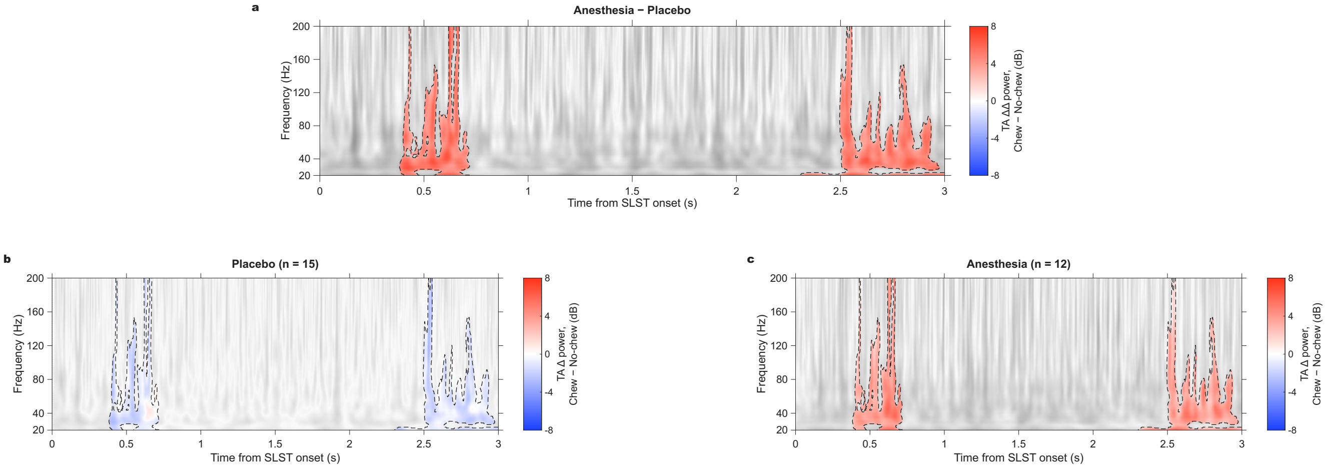


**Figure S1. SLST-onset-aligned sensitivity analysis of the tibialis anterior time–frequency contrast between anesthesia and placebo.** This post hoc analysis evaluated the same between-group difference in Chew minus No-chew power changes as the instruction-locked analysis (Fig. 2), but aligned each trial to its own SLST onset before averaging within participant and block. The analysis included 781 valid TA trials from 27 participants. Power retained the original reference interval of $-1.0$ to $-0.4$ s before the instruction, not before SLST onset; the experimental Baseline block was not used. **a**, Difference between mean participant-level power changes, calculated as $[(\text{Chew} - \text{No-chew})_{\text{anesthesia}} - (\text{Chew} - \text{No-chew})_{\text{placebo}}]$ ($\Delta\Delta$). **b,c**, Descriptive Chew minus No-chew power changes ($\Delta$) in placebo ($n = 15$) and anesthesia ($n = 12$), respectively. Color is retained within the two observed between-group clusters with the greatest mass; the remaining map is displayed in grayscale. The early cluster (C1) extended from 0.383 to 0.719 s after SLST onset ($P_{\text{FWER}} = 0.0866$), and the late cluster (C2) from 2.303 to 3.000 s ($P_{\text{FWER}} = 0.0694$). Both nonrectangular masks reached frequencies between 20 and 200 Hz, without occupying their entire enclosing time–frequency rectangles. Between-group inference used participant group-label permutations with Welch $t$ statistics, 5,000 permutations, a cluster-forming threshold of $|t| > 1.8$, four-neighbor connectivity, and a minimum cluster size of five pixels. Cluster mass summed the magnitudes of same-sign $t$ statistics. Permutation probabilities used the plus-one correction and the maximum cluster mass across both tails over the complete 20–200-Hz by 0–3-s search domain. Neither cluster met the two-sided FWER threshold of 0.05. Correction applied within this test, not jointly across the six muscle-by-contrast tests or analysis versions. Dashed contours identify these nonsignificant observed masks. The same masks are displayed in all three panels; contours in **b,c** therefore do not indicate independently significant within-group clusters. All panels use a common $-8$ to $+8$ dB color scale. Positive values in **a** indicate a greater Chew minus No-chew power change in anesthesia than placebo, not a separately established increase within either group. The group maps descriptively show predominantly negative changes in placebo and positive changes in anesthesia within the shared masks. Although this direction is compatible with the instruction-locked pattern, the onset-aligned analysis did not provide FWER-significant confirmation. Cluster boundaries describe the observed masks, not statistically established effect onsets, offsets, or individual significant pixels; the late cluster reaches the end of the analyzed 3-s window. This sensitivity analysis used the same cohort and does not constitute independent replication. No-chew and Chew denote the assigned oral intervention without and with rhythmic mastication, respectively. TA, tibialis anterior; SLST, Single Leg Stand Test; FWER, family-wise error rate.

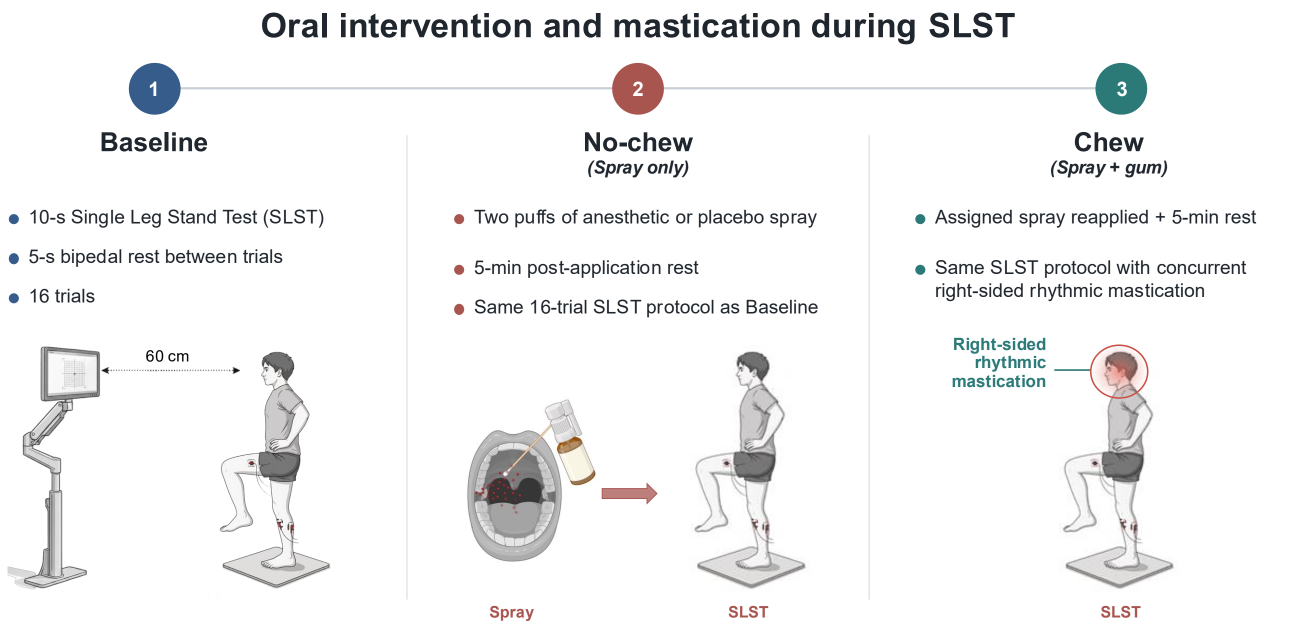


**Figure S2. Experimental paradigm.** Participants completed three fixed-sequence blocks of the Single Leg Stand Test (SLST). The Baseline block comprised single-leg stance without oral intervention. Before the No-chew block, two puffs of anesthetic or placebo spray were administered, followed by 5 min of rest and the same SLST protocol. Before the Chew block, the assigned spray was reapplied, followed by the same 5-min rest period; participants then performed the SLST with concurrent right-sided rhythmic mastication. Each block comprised sixteen 10-s trials separated by 5 s of bipedal rest. Participants viewed a monitor positioned approximately 60 cm away.

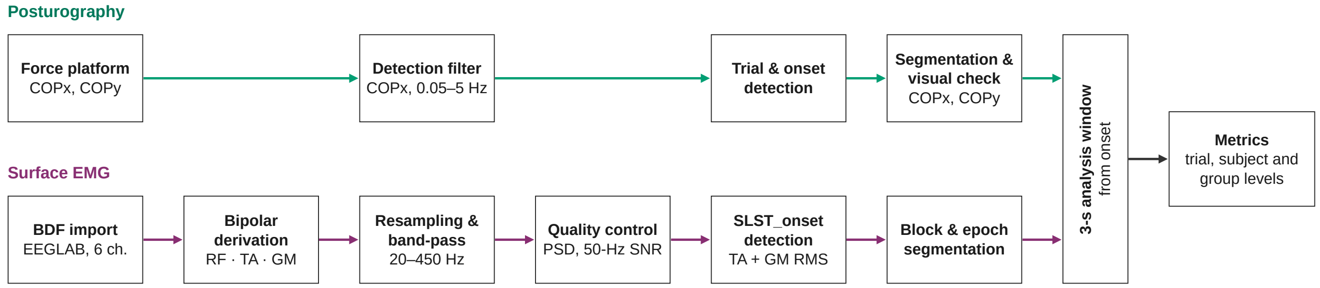


**Figure S3.** **Posturography and surface electromyography processing pipeline.** Posturographic (green) and surface electromyography (sEMG; purple) data were processed in parallel and combined at the trial level (gray). *Posturography:* center-of-pressure (CoP) coordinates in the mediolateral (COPx) and anteroposterior (COPy) directions were derived from force-platform recordings (1,000 Hz). For trial detection only, COPx was band-pass filtered (0.05–5 Hz, second-order zero-phase Butterworth) and smoothed (100-ms moving average); candidate 10-s single-leg stance segments were identified and their onset was refined using hysteresis and moving-variability criteria. Segments were extracted from both CoP coordinates, verified visually for each block, and indexed by participant, block, and trial; outcome metrics were computed from the unfiltered CoP. *sEMG:* recordings from tibialis anterior (TA) and medial gastrocnemius (GM) on the support limb and rectus femoris (RF; auxiliary signal) on the raised limb were imported into EEGLAB, converted from six monopolar channels to one bipolar derivation per muscle, resampled to 1,024 Hz, demeaned, and band-pass filtered (20–450 Hz, fourth-order Butterworth applied forward and backward). Channel quality was assessed from the power spectral density (Welch's method), 50-Hz power-line contamination, and kurtosis. The functional onset of single-leg stance (SLST onset) was detected in each trial from 50-ms RMS envelopes as the first concurrent TA–GM activation exceeding pre-instruction thresholds for at least 150 ms, with a TA-only criterion as fallback; continuous blocks (Baseline, No-chew, and Chew) were then segmented into SLST-onset-aligned trial epochs. Posturographic outcomes used the first 3 s from the CoP-based onset. sEMG amplitude used 0–3 s after SLST onset, whereas coactivation used 0.2–3.2 s after that marker. The streams were organized hierarchically by participant, block, and original trial number for integrated analyses.

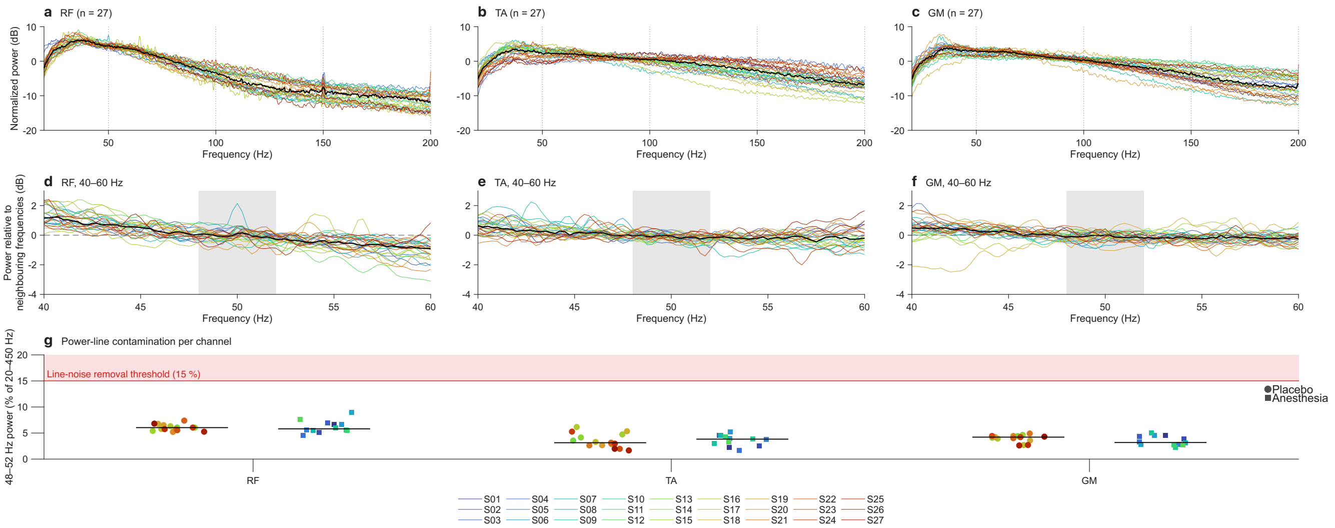


**Figure S4.** **sEMG power-line quality control. a–c**, Power spectra of each participant's bipolar rectus femoris (RF), tibialis anterior (TA), and medial gastrocnemius (GM) recordings (colored lines; black, median), normalized to each channel's mean power between 20 and 200 Hz; dotted lines mark 50 Hz and its harmonics. **d–f**, Detail of 40–60 Hz, normalized to the power of neighboring frequencies (44–47 and 53–56 Hz); a line-noise component would appear as a peak above 0 dB within the shaded 48–52-Hz band. **g**, Proportion of 48–52-Hz power relative to the 20–450-Hz band for each channel (circles, placebo; squares, anesthesia; colors identify participants consistently across panels; black lines, group medians). The shaded area marks the pre-specified 15% threshold for line-noise removal, which no channel reached. RF was recorded as an auxiliary signal and was not analyzed as an outcome.
